# Parsing digital or analogue TCR performance through piconewton forces

**DOI:** 10.1101/2023.11.29.568292

**Authors:** Aoi Akitsu, Eiji Kobayashi, Yinnian Feng, Hannah M. Stephens, Kristine N. Brazin, Daniel J. Masi, Evan H. Kirpatrick, Robert J. Mallis, Jonathan S. Duke-Cohan, Matthew A. Booker, Vincenzo Cinella, William W. Feng, Elizabeth L. Holliday, Jonathan J. Lee, Katarzyna J. Zienkiewicz, Michael Y. Tolstorukov, Wonmuk Hwang, Matthew J. Lang, Ellis L. Reinherz

**Affiliations:** Laboratory of Immunobiology, Dana-Farber Cancer Institute; Boston, MA 02115, USA.; Department of Medical Oncology, Dana-Farber Cancer Institute; Boston, MA 02115, USA.; Department of Medicine, Harvard Medical School; Boston, MA 02115, USA; Department of Chemical and Biomolecular Engineering, Vanderbilt University; Nashville, TN 37212, USA; Department of Dermatology, Harvard Medical School; Boston, MA 02115, USA.; Department of Informatics and Analytics, Dana-Farber Cancer Institute; Boston, MA 02115, USA; Departments of Biomedical Engineering, Materials Science & Engineering, Physics & Astronomy, Texas A&M University; College Station, TX 77843, USA; Department of Molecular Physiology and Biophysics, Vanderbilt University School of Medicine; Nashville, TN 37232, USA

## Abstract

αβ T-cell receptors (TCRs) recognize aberrant peptides bound to major histocompatibility complex molecules (pMHCs) on unhealthy cells, amplifying specificity and sensitivity through physical load placed on the TCR-pMHC bond during immunosurveillance. To understand this mechanobiology, TCRs stimulated by abundantly and sparsely arrayed epitopes (NP_366-374_ /D^b^ and PA_224-233_/D^b^, respectively) following *in vivo* influenza A virus infection were studied with optical tweezers. While certain NP repertoire CD8 T lymphocytes require many ligands for activation, others are digital, needing just few. Conversely, all PA TCRs perform digitally, exhibiting pronounced bond lifetime increases through sustained, energizing volleys of structural transitioning. Optimal digital performance is superior *in vivo,* correlating with ERK phosphorylation, CD3 loss, and activation marker upregulation *in vitro*. Given neoantigen array paucity, digital TCRs are likely critical for immunotherapies.

**One Sentence Summary:** Quality of ligand recognition in a T-cell repertoire is revealed through application of physical load on clonal T-cell receptor (TCR)-pMHC bonds

## Main Text

The vertebrate immune system is comprised of both innate and adaptive cellular components that protect the host from viruses, microbes, toxins, and cancerous transformations (*1*). Innate immunity is rapid and non-specific while adaptive immunity is delayed but decisive, incorporating exquisite specificity and immunological memory (*2–4*). Adaptive humoral and cellular immunity are mediated through lymphocyte receptors, B-cell receptors (BCRs) and T-cell receptors (TCRs), respectively, which undergo somatic rearrangements of gene segments encoding their variable domains during lymphoid development (*5, 6*). This process creates billions of clonotypic structures with the gamut of unique specificities required to recognize diverse pathogens. Without broad repertoire diversity, infectious agents and cancers would overwhelm the mammalian host, as evidenced by pathological sequelae observed in patients with immunodeficiency states (*7*). In contrast to BCRs, αβTCRs are exclusively membrane-bound, lack affinity maturation, and manifest weak monomeric 1-200 µM affinities (*8, 9*). Ligands recognized by BCRs and secreted immunoglobulins are foreign in nature, such as viral envelope proteins. On the other hand, each αβTCR recognizes a foreign peptide bound to a self-MHC molecule, collectively referred to as a foreign pMHC (*5, 10–14*). Foreign pMHCs are arrayed on the surface of a diseased cell or professional antigen presenting cell (APC) at a relatively low copy number amongst a sea of ∼100,000 diverse self-pMHCs.

Given their weak affinities, the strict specificity and sensitivity performance requirements of αβTCRs necessary for cytolytic T lymphocytes (CTLs) to eliminate abnormal cells expressing foreign pMHC were enigmatic. Recent studies solved this paradox by revealing that αβTCRs are force-responsive biomolecules, i.e., mechanosensors that, unlike antibodies, function outside of thermal equilibrium (*15–25*). Tensile forces applied to TCR-pMHC bonds increase their lifetimes and are referred to as catch bonds*. In vivo,* piconewton (pN) forces are placed on an individual αβTCR-pMHC bond through cell motions arising between a T lymphocyte and a target APC during immune surveillance (*15, 16*). The physical load induces conformational changes in the αβTCR heterodimer, reversibly going from a compact to an elongated state, potentially delivering energy to facilitate signaling through perturbation of vicinal membrane lipids and exposure of immunoreceptor tyrosine-based activation motifs (ITAMs) in the cytoplasmic tail of the αβTCR CD3 signaling subunits [(*20*) and references therein].

It follows that αβTCR performance without optimal mechanical load may not accurately reflect biological function *in vivo*, degrading ligand specificity (*17*). Notably, analysis of TCR function *in vitro* is routinely performed at present without such considerations (*26, 27*). Here we use optical tweezers (OT)-based methods to apply the equivalent of biologically relevant pN load to individual TCR-pMHC bonds, revealing differential performance of TCRs recognizing the same pMHC ligand. We discovered a force-driven “molecular resonant” state lasting minutes and involving rapid structural transitioning for those TCRs with the best performance. The value of physiological load application and biophysical parameterization in comparison to immunological metrics like functional avidity (*26–28*) or TCR sequence distance (*29*) measures becomes clear. We posit that those dynamic features of an αβTCR can be linked to facile biomarkers of adaptive immune recognition performance that will track with protective immunity in a clinically useful manner.

### A pipeline of IAV-specific TCRs

To identify αβTCRs directed at two immunodominant but differentially arrayed IAV-specific epitopes, NP_366-374_/D^b^ and PA_224-233_/D^b^, tetramers were used to concurrently isolate CD8 T lymphocytes from lung parenchyma five days post-secondary infection using single cell sorting, RT-PCR molecular cloning, and DNA sequencing (figs. S1A-B). Of 21 MHC-bound epitopes physically identified by mass spectrometry analysis, the copy number on infected cells for NP_366-374_/D^b^ is the highest (> 1,000), while PA_224-233_/D^b^ is amongst the lowest (<10) (fig. S1C) (*30*). Fig. 1A shows a representation of the TCRα (TRA) and TCRβ (TRB) clonotypes with full TRV, TRJ, and CDR3 information provided in data S1. The three most prevalent clonotypes with paired TRA and TRB were used for functional analysis below. NP34 and NP63 utilize the same Vα and Vβ gene segments, differing by only a single amino acid in Vα CDR3 (Fig. 1A). By contrast, NP41 utilizes entirely different V gene segments encoding a divergent VαVβ recognition module.

**Fig. 1.**
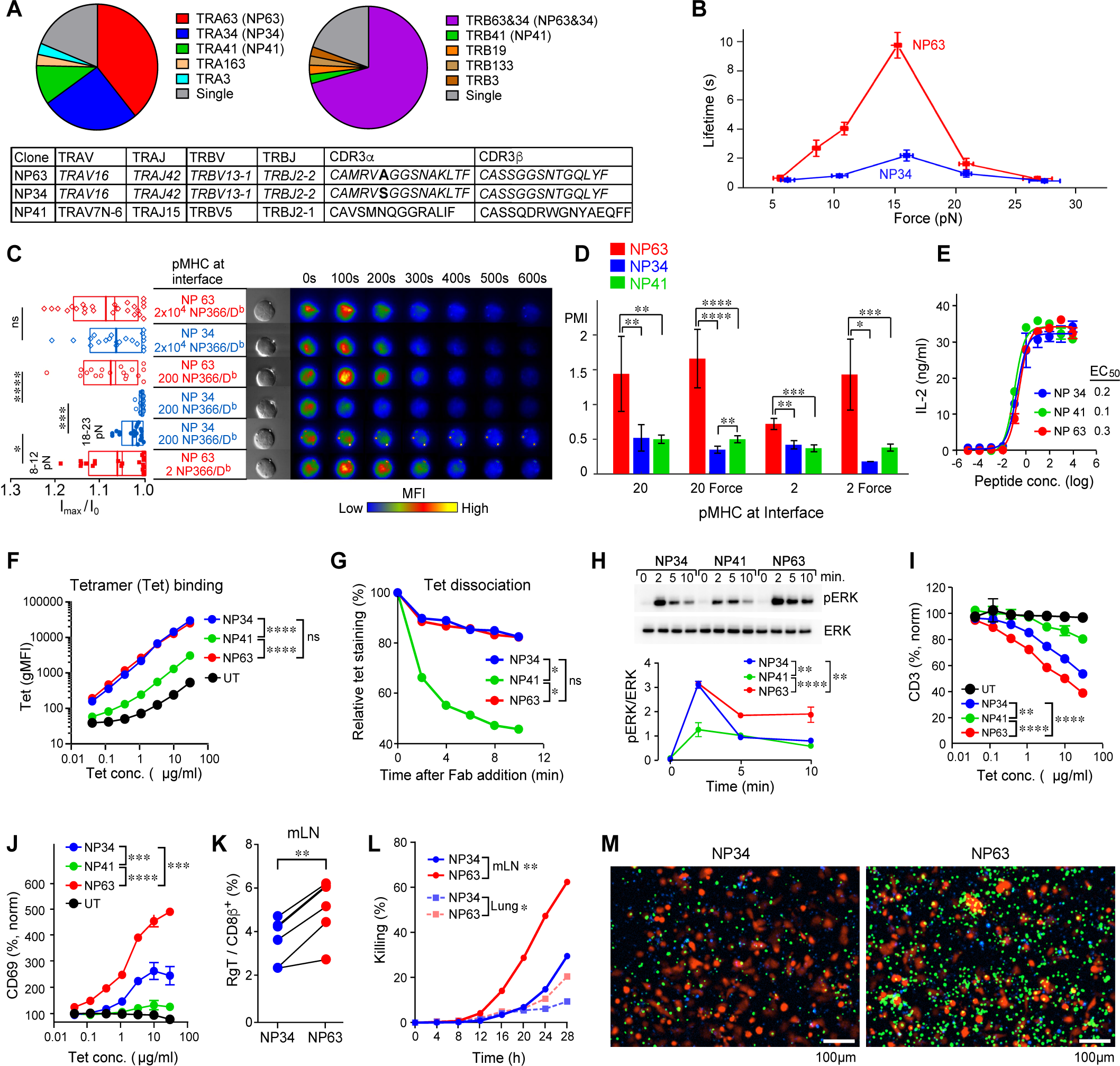
TCRs recognizing a high copy number pMHC array on IAV infected cells function in either an analogue or digital performance mode. **(A)** Repertoire analysis of NP_366-374_/D^b^-specific TCR. Pie charts show the frequency of individual CDR3α (left) and CDR3β (right) clonotypes, with 2 or more cells colored as indicated, and single-cell clonotypes in gray. NP34, NP63, and NP41 TCRs utilize the TRV, TRJ, and CDR3 sequences summarized in the bottom table. NP34 and NP63 share the same CDR3 sequence (italic letters) except for letters in boldface. (**B)** SMSC assay to measure bond lifetime versus force for NP34 and NP63 TCR transfectants generated by retroviral transduction into BW5147CD8αβ^+^TCRαβ^-^ parental cells. Data are shown as mean ± SEM. Peak lifetimes occur at 15pN, 10 s and at 16 pN, 2 s for NP63 and NP34 respectively. **(C)** SCAR assay to measure the magnitude and the longevity of calcium flux in NP34- and NP63-BW cells using either 2, 200, and 20,000 NP_366-374_/D^b^ copy numbers at the bead–cell interface with or without an optimal vectorial force. The calcium flux signal is indicated as the ratio of maximum fluorescence intensity (I_max_) to the initial fluorescence intensity (I_0_) of the Ca^2+^-sensitive dye. Each dot in the plot represents a single-cell experiment. The rectangles span SD with mean and median values shown as thick and thin lines, respectively. Pictures to the right are representative DIC (Differential Interference Contrast) images of a cell-bead pair in SCAR experiment, and time-lapse images of intracellular free Ca^2+^ release for representative cells. MFI, mean fluorescence intensity. (**D)** Quantification of PMI (Predicted Mean Intensity; the average signal level for triggered cells multiplied by the percentage of cells that trigger in each category) for NP34, NP41, and NP63 with the indicated interfacial number of pMHC with or without force application. These data were collected separately from **C**, using a different dual fluorescence OT system with greater fluorescence sensitivity (see SCAR section in Materials and Methods). **(E)** IL-2 production from the indicated TCRαβ-transduced BW cells after stimulation with titrated NP_366-374_ peptide presented on R8 APCs. Peptide concentrations are in ng/ml where Log_10_ values are shown on the x-axis (0: 1 ng/ml). (**F**) NP_366-374_/D^b^ tetramer binding (geometric mean fluorescence intensity, gMFI) to TCR-transfected or untransfected (UT) BW5147CD8αβ^+^ cells following overnight treatment at 37 °C to trigger T cell activation in **I** and **J**. Fluorescence intensity was measured by flow cytometry. **(G)** Tetramer dissociation assay. Indicated cells were incubated with WT tetramer and then treated with the 28-14-8 antibody Fab fragment against H-2D^b^/H-2L^d^ at indicated time points. (**H)** Western blot analysis of ERK phosphorylation (p-ERK) in cells after NP_366-374_/D^b^ tetramer stimulation. Blots on top show one of three representative results with the p-ERK to ERK ratio at each time point after activation is shown below. (**I, J**) Loss of surface CD3 and concomitant upregulation of CD69 expression after stimulation shown in (**F)**. The fluorescence intensity of CD3 (**I**) and CD69 (**J**) was measured by flow cytometry and normalized using the non-tetramer stimulated value. **(K)** Frequency of NP34 and NP63 Rg T cells in mediastinal LN (mLN) of mixed RgC mice 7 days after PR8 IAV infection. **(L)**, Quantification of real-time Rg T cell-mediated killing of PR8-infected LET1 cells over time (h, hours). NP34 and 63 Rg T cells were derived from mLN or lung of RgC mice (dpi 7). (**M**) Representative images of the killing assay for mLN Rg T cells at 20 hours in **L**. LET1 cells were visualized by transduced mCherry, apoptosis is in green, and Rg T cells are in blue. For **D**, data are shown as mean ± SD. For **E-G,** and **I-M**, data are representative of 2-4 independent experiments and are shown as means ± SDs of technical replicates (**E, F, I,** and **J**) and of 6 mice (**K**). For **H**, data are shown as means ± SDs of 3 independent experiments. Some error bars are invisible due to small SDs (**E, F, H, I,** and **J**). For all data with statistics, ****P <0.0001, ***P<0.001, **P<0.01, *P<0.05; ns, not significant. P values were calculated by one-way ANOVA (**C** and **D**), comparing slopes of linear regression (**F, I, J,** and **I**), by the Kolmogorov-Smirnov test (**G**), by regression using trend line analysis models accounting for interexperimental variability (**H**), or by paired t-test (**K**).

### TCR recognition of dense pMHC under load

Given the near identity of the TCR sequences directed at the same pMHC, we assumed that NP63 and NP34 would yield a similar functional profile. To our surprise, however, clear distinctions emerged. To rule out differences in TCR copy number, adhesion molecules and/or signaling pathways leading to divergent functional outcomes, each TCR was retrovirally transduced into the same parental CD8αβ^+^ TCR^-^ BW5147 recipient cell line and selected for comparable TCRαβ expression (fig. S2A). Two key OT-based assays were used to interrogate transfectants (figs. 2B-C). Single-molecule single-cell (SMSC) measurements tether pMHC molecules to a bead through a DNA rope and present the bead to a surface of a coverslip-bound T cell. As the tethered bead is pulled away, it is displaced from the trap center and is held until the TCR/pMHC bond is broken, revealing the bond lifetime for a given force. Tether formation probability as well as peak lifetime, peak force, and width of catch bonds can also be determined. In the single cell activation requirement (SCAR) assay a pMHC-coated bead is trapped and moved to form interfacial contact with a T cell containing a fluorescence-based reporter of intracellular calcium concentration. An extended calcium flux in these cells is used as an indicator of T cell activation. Beads of varying pMHC densities can be used to judge TCR performance quality and determine the interfacial pMHC copy number required for activation, either with or without force. The tug by the OT mimics load between a T cell and an APC (or an infected cell) exerted through their respective actomyosin machineries (*31*).

Fig. 1B demonstrates catch bond profiles. Bond lifetimes for both NP63 and NP34 TCRs peaked at ∼15 pN, but NP63 had a 5-fold longer bond lifetime than that of NP34. NP41 had a bond lifetime equivalent to NP34 (fig. S2D), yet its low tethering probability (fig. S2E) required measurement using a single molecule (SM) assay illustrated in Fig. 3A as opposed to SMSC. The bond lifetime difference between NP63 and NP34 is further reflected by their differential responsiveness in the SCAR assay (Fig. 1C). Although both induced calcium flux with similar kinetics at high bead copy number (20,000 interfacial pMHCs per bead), only NP63 could be activated by 200 pMHCs arrayed per bead without load. Even 18-23 pN force application did not cause many NP34 T cells to induce calcium flux in this system. In contrast, at 8-10 pN, NP63 was readily activated with as little as two pMHCs per bead, the lowest interfacial ligand concentration achieved in this study.

To further elucidate activation thresholds, we adapted the SCAR assay to a microscope specifically designed for SM fluorescence detection coupled with a sensitive camera temporally gated with excitation at very low illumination levels. These conditions virtually eliminated photobleaching and extended facile monitoring of calcium activation beyond 10 minutes. We thereby assayed the activation profiles more thoroughly for NP63, NP34 and NP41 at 20 and 2 interfacial copy numbers with and without force. While every condition revealed an activation capability, the greatest impact observed was on the percentage of activated cells. (fig. S2F). This percentage combined with the activating signal level yields the predicted mean intensity (PMI), which revealed a statistically greater performance by NP63 (Fig. 1D, fig. S2G). We refer to NP63 TCR responsiveness as digital since only a couple of pMHC ligands are required for activation. In contrast, the NP34 TCR responsiveness is termed analogue, requiring multiple pMHC ligands to stimulate the T cell in a graded manner.

### Immunological assay comparisons

Functional avidity measurements using cytokine production as a readout are commonly performed to assess TCR quality through examination of T cell responsiveness to APCs cultured overnight with different peptide concentrations (*28*). The assay is technically straightforward, but its interpretation is complex, given myriad cellular components involved including adhesion molecules, co-receptors and kinases affecting cytokine secretion. As shown in Fig. 1E, the functional avidity of all three NP TCRs is comparable. While NP_366-374_ /D^b^ tetramer binding and dissociation were comparable for NP34 and NP63 (Fig. 1F and G, respectively), the binding to NP41 was the weakest and manifested the fastest dissociation rate. The wild-type (WT) NP_366-374_/D^b^ tetramer (Tet) was used for this dissociation assay because NP41 does not interact with a CD8 binding-site (BS) mutant (Mut) MHC molecule (fig. S3A). Collectively, these findings reveal that digital vs analogue performance amongst TCRs cannot be discerned by the commonly used metrics of functional avidity or pMHC tetramer binding or dissociation.

Nevertheless, since tetramers can mediate crosslinking of adjacent TCR ectodomains on a T cell, themselves tethered internally to the actin cytoskeleton, we reasoned that mechanical force applied following such *in vitro* exposure could be used to interrogate downstream activation features. We tested whether differential mechanosensing amongst TCRs elicits divergent signaling responses. As the binding profiles of NP_366-374_ /D^b^ tetramer for NP63, NP34 and NP41 were similar at 37°C (Fig. 1F) and at 20°C (fig. S3A), T cell activation at 37°C following tetramer binding could be readily studied. Rapid phosphorylation of ERK (pERK) was detected within 2 minutes, where the greatest magnitude and persistence were observed for NP63 (Fig. 1H, and data S2). Furthermore, the same binding leads to a differentially graded CD3ε surface loss among the three TCRs (Fig. 1I) and distinctive upregulation of the early C-type lectin activation marker CD69 (Fig. 1J, and fig. S3B).

As these studies involved *in vitro* assays, we next determined whether *in vivo* activation of NP63 and NP34 T cells differed upon IAV infection. To this end, we created single or mixed retrogenic T cell (Rg-T) mice bearing each TCR expressing T cell alone or together using FACS sorting of naïve retrogenic CD8^+^CD44^-^ T cells for those adoptive transfer experiments into B6 mice followed by IAV infection, as explained later. We quantified the mediastinal lymph node (mLN) representations at day 7 post infection. As shown, NP63 CD8^+^ T cells expanded significantly more than NP34 in the mixed retrogenic setting and revealed the greatest incorporation of EdU, the nucleoside analogue of thymidine, into DNA during the S-phase of cell cycle (Fig. 1K and figs. S4A-C). In addition, when Rg-T cells were sorted and tested for cytolytic activity against IAV-infected mCherry^+^ LET1 type I pneumocytes, as monitored continuously over 28 hrs *ex vivo,* mLN NP63 were faster and better at killing than were NP34 T cells (Figs. 1L-M). This was also the case for lung-derived NP63 T cells (figs. S4D-E). The efficacy of IAV infectious doses that supports T-cell mediated killing of LET1 cells roughly correlates with intracellular LET1 NP expression by FACS analysis (fig. S4F). As shown in figs. S4G-I, Rg-T cell numbers in lung and the level of viral titer reduction at day 7 post-IAV infection were comparable for NP34 and NP63, consistent with the notion that the high copy number of the NP_366-374_ /D^b^ complexes allows both digital and analogue TCR performance to be effective.

### Sparse pMHC recognition under load

Corresponding analysis of PA_224-233_/D^b^ -specific TCRs (Fig. 2A and data S1) identified two common TCRs termed PA27 and PA59, and a less frequent PA25. They differ in sequence aside from PA25 and PA59 that share a Vβ gene segment and very similar CDR3β. The SMSC force vs. bond lifetime analysis revealed that all three TCRs exhibited catch bond behavior, like those of the NP series (Fig. 2B and Fig. 1B). However, the PA59 maximal bond lifetime (75 s) was longer, and it occurred at a significantly greater force, 21 pN. Nonetheless, all three PA TCRs manifested digital performance, being triggered by 2 PA_224-233_/D^b^ molecules per bead-cell interface in the SCAR assay (Fig. 2C). While PA27 and PA25 triggered well in the 8-12 pN range, PA59 required a higher force, i.e., 16-18 pN, which is consistent with the SMSC result (Fig. 2B). The duration of Ca^2+^ flux was longer for PA27, shortest for PA25, and with the greatest intensity for PA59 at high force (Fig. 2C, fig. S5, and movies S1-S3). Paradoxically, the functional avidity assay indicated that PA27 was the weakest TCR based on EC_50_ (Fig. 2D). Of note, the IL-2Rα expression (CD25) on the BW cell lines after peptide stimulation is comparable (fig. S6A). Hence, IL-2 depletion is not responsible for the discrepancy.

**Fig. 2.**
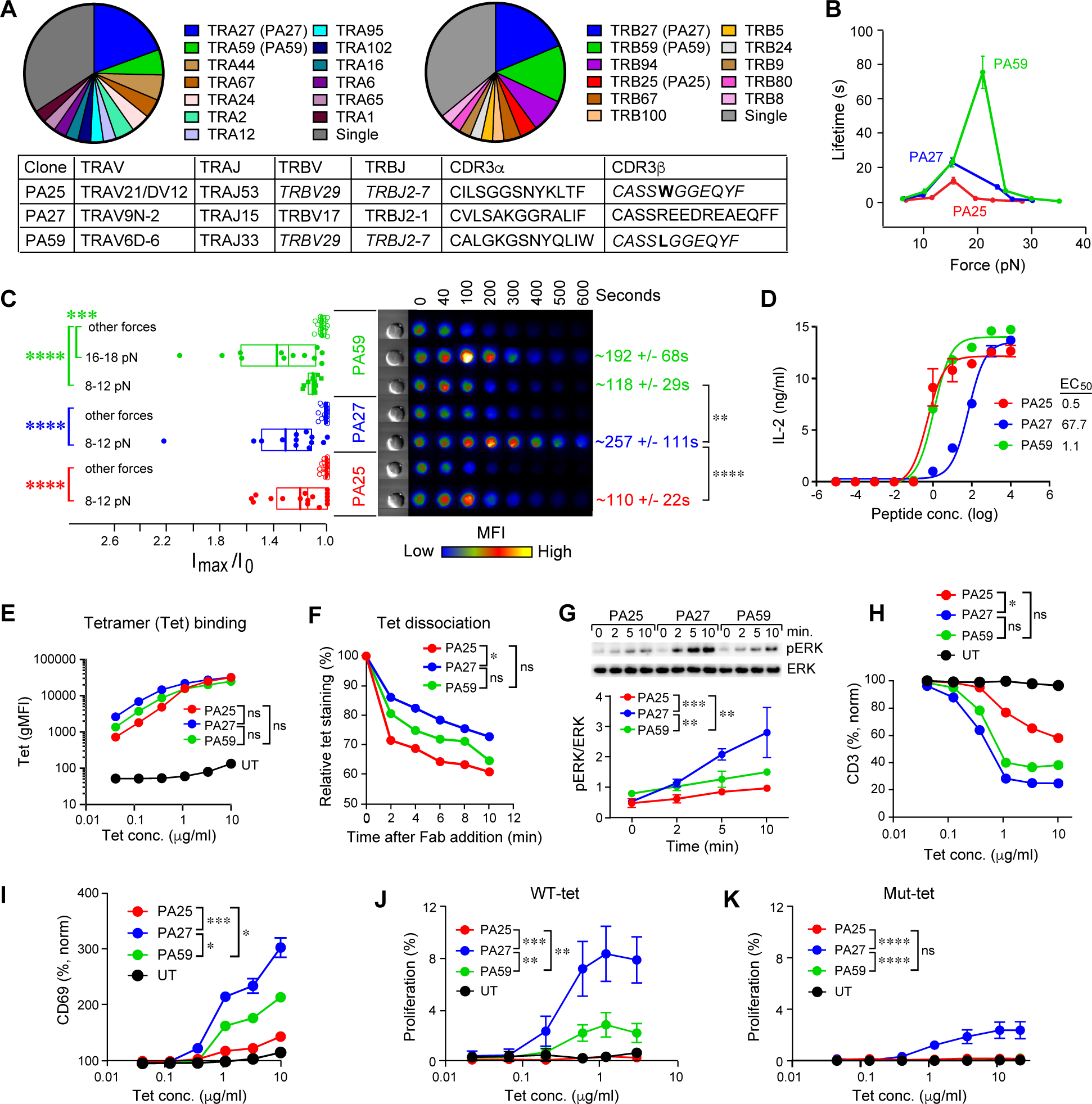
A sparse immunodominant pMHC array exclusively elicits TCRs with digital performance but distinguishable activation features. **(A)** Repertoire analysis of PA_224-233_/D^b^-specific TCRs. Pie charts and the table are as explained in Fig.1. (**B**) SMSC measurement of bond lifetime versus force for PA25, PA27, and PA59. Peaks of the catch bond curves occurred at 21 pN and 75 s for PA59, and at 15 pN at 23 s and 13 s for PA27 and PA25, respectively. Data are in mean ± SEM. (**C)** SCAR assay of indicated transductant with all three PA TCRs triggered by two PA_224-233_/D^b^ molecules on a bead in conjunction with external force. Other forces indicate outside the optimal range, < 8 pN and > 12 pN for PA25 and PA27, and < 8 pN and > 18 pN for PA59. See Fig. 1C for explanation of plots and images. (**D)** IL-2 assay for indicated BW transductants after stimulation with titrated PA_224-233_ peptide presented on R8 cells as explained in Fig. 1E. (**E)**, Tetramer binding measured after overnight incubation with WT PA_224-233_/D^b^ tetramer treatment. The fluorescence intensity was measured by flow cytometry. **(F)** Tetramer dissociation assay completed after PA25-, PA27-, and PA59-BW cells were incubated with a PA_224-233_/D^b^ tetramer harboring a CD8BS-MHCⅠ mutation and then treated with 28-14-8 antibody Fab fragment for the indicated time point before FACS analysis. (**G)** Western blot analysis of p-ERK as described in Fig. 1H. **(H, I**) Change of surface CD3 (**H**) and CD69 (**I**) expression with increasing concentration of WT tetramer as shown in **E**. Indicated BW cells were incubated at 37 °C overnight with tetramer, then fluorescence intensity of CD3 (**H**) and CD69 (**I**) was measured by flow cytometry and analyzed as in Fig. 1I and J. **(J, K)** Differential proliferation of BW transductants to WT (**J**) and CD8BS-mutant (**K**) tetramers. Measurements were obtained 30 minutes after addition of WT (**J**) or mutant (**K**) tetramers. The frequency of proliferation was determined by FSC-A vs. SSC-A plot by flow cytometry and normalized by the non-tetramer stimulation value. For **D-F** and **H-K**, data are representative of 2-4 independent experiments and are shown as mean ± SD (**D, E, H,** and **I**) and ± SEM (**J** and **K**) of replicates. For **G**, data are shown as mean ± SD of 3 independent experiments. Some error bars are invisible due to small SDs or SEMs (**D, E, G-K**). For all data with statistics, ****P <0.0001, ***P<0.001, **P<0.01, *P<0.05; ns, not significant. P values were calculated by one-way ANOVA (**C**), comparing slopes of linear regression (**E, H**-**K**), by the Kolmogorov-Smirnov test (**F**), and by regression using trend line analysis models accounting for interexperimental variability (**G**).

The above results suggest that digital PA receptors are not monolithic, but rather function in distinct ways under force. Nevertheless, WT-tetramer binding assays revealed no substantial differences among these TCRs (Fig. 2E and fig. S6B) while CD8BS-mutated tetramer binding was weaker only for PA25 (fig. S6B). PA27 and PA59 tetramer dissociation were similar, but both were at a slower rate than PA25 (Fig. 2F). On the other hand, WT-PA_224-233_/D^b^ -tetramer activation assays showed that PA27 manifested prolonged Erk-mediated activation (Fig. 2G and data S2). CD3ε downregulation was also the largest for PA27 but not statistically distinguishable from PA59 (Fig. 2H and fig. S6C). Differential CD3 surface loss increased at higher temperature but with almost no change in tetramer binding, suggesting an active cellular mechanism of CD3 downregulation likely through tetramer stimulation fostering kinase-dependent internalization (*32, 33*). and/or dissociation of CD3 dimers from TCRαβ (*20*) (fig. S6D). PA27 also showed the greatest differential CD69 upregulation (Fig. 2I and fig. S6C), and the greatest proliferation response to WT-as well as CD8BS mutated-PA_224-233_/D^b^ tetramers (Figs. 2J-K and figs. S6E-F). Of note, none of the NP TCRs proliferated following CD8BS-mutated NP_366-374_ /D^b^ tetramer stimulation (fig. S6G).

### SM OT analysis of digital TCRs

We next evaluated the performance of the digital PA TCRs using SM analysis. We adapted our SM assay to a geometry where the TCR is bound to a 1.23 µm bead through the anti-leucine zipper mAb 2H11 and pMHC is bound to a second 1.23 µm bead via a DNA linkage (Fig. 3A). The SM “dual bead” assay (SM_db_) was performed by positioning the two traps to initiate tether formation followed by pulling the linkage taught to a predetermined force window. This assay was performed on a LUMICKS m-Trap microscope where microfluidic flow introduces beads to populate the traps and a measurement routine calibrates the system.

**Fig. 3.**
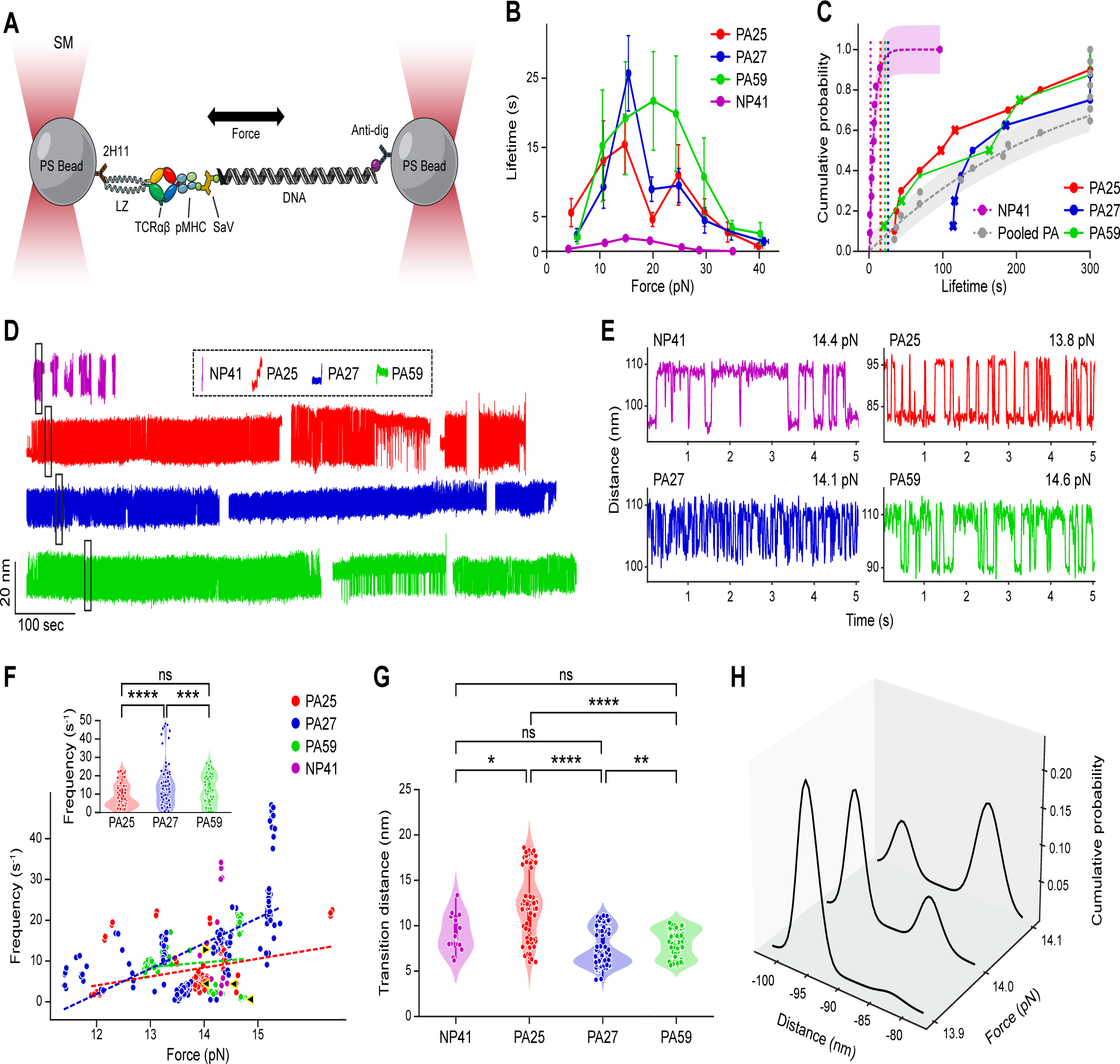
A single molecule dual bead OT system discriminating mechanosensing performance of digital TCRs. **(A**) SM_db_ system cartoon. **(B)** Bond lifetime vs. force for PA25 (red, n= 175), PA27 (blue, n=237), PA59 (green, n=192), and NP41 (magenta, n=57). Color convention is maintained throughout Fig. 3. Lifetimes are binned every 5 pN and are plotted as mean bond lifetime ± SEM. (**C)** Cumulative probability plot for continuous volleying segments. In comparison, the peak lifetime from the catch bond curves for each clone is noted by a vertical dashed line with the corresponding color. Symbols depicted by an X indicate termination by the user. PA25 (red, n= 10), PA27 (blue, n=8), PA59 (green, n=8), and NP41 (magenta, n=11). PA events were pooled (grey) and fit to the function 𝑦 = (1 − 𝑒^−𝑥⁄𝑡^). 95% confidence intervals are denoted by magenta (NP41) and grey (pooled PA) shaded areas. **(D**) Representative traces of continuous volleying segments. Multiple extended hopping traces are shown for each clone, separated by a blank space. All traces shown are in 13.8 – 14.5 pN force range. Sample traces from each catch bond curve in the same force range that did not show reversible transitions are shown in the dotted box. **(E**) Zoomed in sections of the black rectangles in Fig. 3D. **(F)** Hopping frequency vs. force in 10-s segments. Markers for NP41 (magenta open circles) represent segments shorter than 10 seconds due to the lack of sustained volleying. Note that the force indicated is the force applied to the folded state and the force upon opening is 1-1.5 pN smaller. Dashed lines are linear regressions used to guide the eye. Arrowheads with yellow borders indicate elements used in Fig. 3E. Violin plots in the top left insert show pooled frequency for PA’s. (**G**) Transition distance for 10-s segments. The range of the transition is indicated. We consider the major transitions between the distributions of the two major dwell states, excluding small transitions < 5 nm observed within a particular state. **(H**) Example of SM position distributions vs force for the PA25 sample segment in Fig. 3D showing shift in population between two major states as a function of force near the critical force of 14 pN. For all data with statistics, ****P <0.0001, ***P<0.001, **P<0.01, *P<0.05; ns, not significant. Fits and 95% confidence ranges from the fit parameters are shown as dashed lines and shaded regions, respectively (**C**). P values were calculated by Kruskal-Wallis tests (**F insert and G**).

PA27 and PA25 showed strong catch bond peaks around 15 pN, while PA59 showed a much broader distribution of lifetimes and broader peak force range. PA27 displayed the longest and narrowest peak lifetime (Fig. 3B). The wide lifetime spread, attributed to an increase in instrument response time capturing short lived interactions for the SM_db_ assay, and variation in curve shape relative to SMSC (Fig. 2B) prompted a closer look at the lifetime distribution within each force window. A common observation was that the cumulative probability of lifetimes within each bin largely fit a double exponential model, wherein there is one time constant for quick dissociation (< 2 sec) and another for an extended lifetime (fig. S7A). The peak bin of ∼15 pN spanning the critical force for conformational transition, was significantly higher, demonstrating a ∼45-50% increase in lifetime for PA27 relative to PA25 and PA59. Fit parameters converged to 29.4 +/- 5.3 s and 28.2 +/- 3.7 s for PA25 and PA59, respectively, compared with 42.6 +/- 4.4 s for PA27 (fig. S7B). Overall, PA clones have longer bond lifetimes compared to NP41-NP_366-374_ /D^b^ which exhibited a much shorter catch bond peak lifetime and more difficulty in initiating tether formation in the same SM_db_ assay (Fig. 3B).

In the SM_db_ assay, force can be incrementally altered during a measurement by slight adjustment in the trap separation. By actively maintaining tethers at or near a critical force, we were able to observe repeat reversible transitioning between extended and compact states with a corresponding extension of bond lifetime, a state we refer to as volleying. During volleying near the critical force, the linkage persisted for several minutes and in some cases more than 5 minutes. To illustrate, we plot the cumulative probability of volleying lifetimes, which are much longer than the catch bond lifetimes (vertical dashes in Fig. 3C). The population of volleying segments at the 5-minute lifetime mark for digital PA27, PA59, and PA25 were 25%, 12.5% and 10%, respectively, although some traces were artificially ruptured by the user as noted (Fig. 3C). In contrast, although NP41 transitioned, it lasted only ∼5 seconds on average (Figs. 3C-E). The cumulative distribution for NP41 fit to a time constant of 6.8 +/- 0.85 seconds. For additional comparison we pooled lifetimes of PA-specific TCRs that either terminated naturally or survived to the 5-minute mark, yielding a time constant of 264 +/- 25 seconds which is far beyond peak catch bond lifetimes and ∼40 fold longer than that of NP41 (Fig. 3C).

Long periods of sustained volleys were divided into 10-s segments for further study. Analysis of the transition frequencies vs. force of these sustained volleys showed that PA25, PA27 and PA59 largely behave similarly (Fig. 3F), but that PA27 has potential to transition faster (Fig. 3F insert) and for a slightly longer period of time (Fig. 3C). NP41 generally transitioned at a low frequency compared to the others at the same force, but all showed a spread of frequency spanning 5-30 Hz (Fig. 3F). Frequencies in Hz for PA27, PA25 and PA59 were 14.1 +/- 10.7, 7.8 +/- 6.1 and 9.5 +/- 7.0 (avg. +/-), respectively (Fig. 3F insert). The average transition distances were similar, ranging from 8-12 nm but with different distributions (Fig. 3G). Note the two distinct PA27 transition distances, for example. All three PA-specific clones had average critical forces for volleying in the 13.7-13.9 pN range, whereas NP41 volleyed on average at 14.3 pN. The critical force for volleying generally centered around ∼14 pN below which the population distribution favored the closed state and above favored the extended state (Fig. 3H).

### *In vivo* transcriptome of digital TCRs

To assess the impact of differential digital TCR performance *in vivo*, we generated Rg mice by transferring *Rag2*^-/-^ hematopoietic stem cells retrovirally transduced with each TCRαβ clonotype and an IRES-linked fluorescence protein into recipient *Rag2*^-/-^ mice (Fig. 4A). Subsequently, an equal number of naïve Rg-T cells from those Rg mice were adoptively transferred into B6 mice (Rg-chimera mice, RgC mice) followed by PR8 infection (Fig. 4A and fig. S8A). FACS analysis allowed for detection of the Rg-T cells using a combination of the fluorescent protein and antibodies against Vβ (GFP^+^ V9β^+^ for PA27, GFP^+^ Vβ7^+^ for PA59, and mCherry^+^Vβ7^+^ for PA25), and revealed that the dominant Rg-T cells in mLN are PA27, followed by PA59, and then by PA25 at 7-days post-infection (dpi 7) (Figs. 4B-C). Those data are consistent with increased *in vivo* EdU incorporation by PA27 relative to PA25 and PA59 that was not significantly different from one another (Fig. 4D and fig. S8B). Of note, percentages of all three Rg T cells relative to CD8β^+^ T cells were equivalent in lung (figs. S8C-D).

**Fig. 4.**
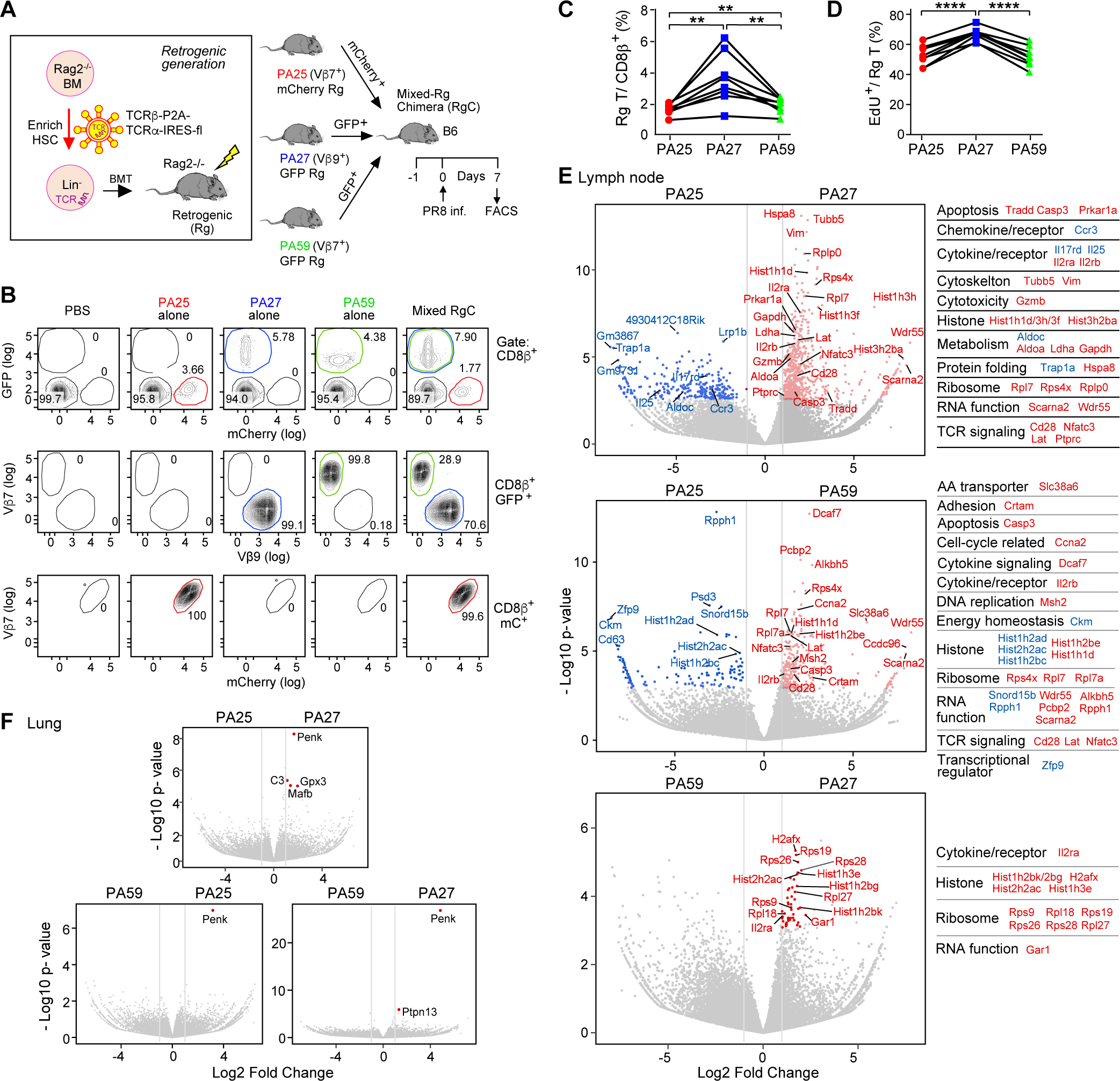
In vivo transcriptomes and expansion of T cells expressing digital TCRs during the acute IAV response. **(A**) Experimental schema to analyze *in vivo* behaviors of three distinct clonotypic PA_224-233_/D^b^ TCRs in the same mice. Rg mice were generated by transferring *Rag2*^-/-^-derived hematopoietic stem cells (HSC) after transduction of retroviruses containing TCRβ-P2A-TCRα with a fluorescence-protein (FP) gene into irradiated *Rag2*^-/-^ mice (left). Subsequently, mixed RgC mice were generated by adoptively transferring an equal number of naïve FP^+^CD8β^+^CD44^-^ T cells from PA25-mCherry, PA27-GFP, and PA59-GFP Rg mice into recipient B6 mice, followed by PR8 infection of the latter 24 hours post-transfer. Rg T cells were analyzed on day 7 post-infection. (**B)** Representative contour plots showing the frequency of Rg T cells in mLN of mixed RgC mice. PA25 T cells were identified as CD8β^+^mCherry^+^Vβ7^+^ (TRBV29^+^) cells (red circle), PA27 T cells as CD8β^+^GFP^+^Vβ9^+^ (TRBV17^+^) (blue circle), and PA59 as CD8β^+^GFP^+^Vβ7^+^ (TRBV29^+^) (green circle) by flow cytometry. (**C)** Quantification of the frequency of Rg T cells in mLN. (**D)** Quantification of the percentage of EdU^+^ Rg T cells. (**E)** Volcano plots of bulk RNA-Seq expression data of FACS isolated clonotypes displaying differentially expressed genes (DEG) as colored dots (red or blue) between PA25 and PA27 Rg T cells (top), PA25 and PA59 (middle), and PA59 and PA27 (bottom) in mLN. Each gene of note is functionally categorized and listed on the right. (**F)** Volcano plots indicating the paucity of DEG between each Rg T cells in lung. For **C** and **D**, data are representative of four independent experiments. P values were calculated by paired t-test. **P < 0.01; ****P <0.0001. For **E** and **F**, DEG (the red and blue dots) are identified as log2 Fold-Change >1 (vertical gray lines) and adjusted P value ≤ 0.05 (Wald test with Benjamini-Hochberg correction).

To exclude the possibility that the binding of mAbs to the TCRs used for cell sorting might have induced TCR signaling and impacted gene expression, we used a third fluorescence protein BFP, for PA27, thus generating an “untouched” TCR labeling and sorting system. Subsequently, we generated mixed RgC mice and performed bulk RNA-seq at 7 dpi (fig. S9A and data S3). Two of the 12 samples, one from PA25 mLN and one PA59 lung were excluded from further analysis due to low RNA quality. Principal component analysis (PCA) shows that each Rg T cell type in mLN is clustered, whereas those in lung are scattered and undistinguishable (fig. S9B). Compared to PA25 and PA59, PA27 T cells differentially upregulate genes (635 and 48 genes, respectively), including those involved in TCR signaling, cytotoxicity, cytokine and chemokine receptors, ribosomes, metabolism, and apoptotic genes (Fig. 4E and data S4). Gene set enrichment analysis (GSEA) also revealed that cell cycle pathways are significantly upregulated in PA27 compared to both PA25 and PA59, and the TCR signaling pathway is additionally upregulated in PA27 T cells compared to PA25 (fig. S9C). Compared to PA25, PA59 T cells upregulate 264 genes, including TCR signaling, ribosome, and cell cycle genes (Fig. 4E), although the GSEA was not statistically significant (fig. S9C).

In contrast to the many differentially expressing genes amongst the three PA-Rg T cells in mLN, there were almost no upregulated genes in those same PA-Rg-T cells in lung, except for a few most prominently displayed in PA27 T cells such as *Penk*, *Mafb, Gpx3 and C3* (Fig. 4F and fig. S9D). We note that the overall comparability of the three types of Rg T cells in lung clearly is not due to their unresponsiveness because all significantly upregulated genes associated with TCR signaling, inflammation, and cytotoxicity compared to those in mLN including *Nur77, Zap70, Nfat, Gzmb, Prf1, Il2ra* (figs. S10A-E). The equivalence of gene expression in the lung is probably a consequence of the collective T cell activation resulting from αβTCR triggering in the context of inflammation with attendant cytokine- and chemokine-mediated signaling spawning modest but equivalent anti-viral activity in those PA-Rg T cells (fig. S10F). Of note, *ex vivo* killing assay shows no PA-Rg T cell-mediated cytotoxicity of LET1 cells (fig. S10G), most likely because of the virtual absence of PA peptides presented on a major fraction of LET1 cells compared to, on average, a 5-10 fold higher number on the DC2.4 dendritic cell line post infection^19^. Thus, it is likely that PA-T cells expand by recognizing PA_224-233_/D^b^ presented on DC in mLN and contribute to virus clearance in lung through production of cytokines and chemokines in a bystander fashion.

## Discussion

Over 200 million years of jawed vertebrate (*Gnathostomata*) evolution, mammals have developed an αβ T-cell lineage immune system that utilizes mechanosensing to detect sparse pMHC ligands. This advancement enhances sensitivity by 1,000-10,000 fold compared to T cells operating without bioforces (*15*). The Cβ FG loop plays a crucial role by stabilizing the Vβ-Cβ domain interaction, controlling pMHC interaction surface orientation and contributing to TCR specificity, sensitivity, and bond lifetime (*17, 25*). Load applied across the TCR-pMHC interface stabilizes interdomain contacts both within the TCRαβ domains and with pMHC, thereby fostering access to transitioning between compact and extended TCRαβ conformations, sustained by external force arising from actomyosin machinery in the T cell and APC (*15, 16*). Unsurprisingly, deletion of the Cβ FG loop dramatically degrades αβTCR-pMHC recognition function (*17*). In contrast to αβT cells, γδT cells lack the equivalent of the elongated Cβ FG loop, as it is apparently unnecessary for their recognition of abundant non-peptidic surface ligands. Consequently, γδT cells manifest neither catch bonds nor structural transitions (*34*). These findings fill a gap in the understanding of early components of TCR-mediated T cell activation previously investigated more broadly (reviewed in ref 16).

Here we show that with proper force feedback, the TCR-pMHC bond can adopt an unprecedented resonant state revealing lifetimes ten-times greater than peak catch bond lifetimes (Fig. 3). Given the fixed separation between traps, which includes the TCR-pMHC bond, DNA linkage, beads and optical springs (with physiologically relevant stiffnesses in the 0.2-0.3 pN/nm range), a sudden increase in length of the TCR-pMHC bond results in transient relaxation and corresponding reduction of force. This, in turn, shifts the energy landscape to just below the equilibrium force favoring transition back to the compact state and a reset of the cycle. Unlike protein unfolding where the distance to the transition state in the forward direction, i.e. unfolding, is very short (in the Å or single nm range) compared to the multiple nm-scale refolding distance and thus making it energetically unlikely to refold with a sustained load, the nm-level forward and reverse TCR-pMHC transition state distances are more balanced. This greater parity fosters the rapid volleying observed within a narrow “resonant window” of the critical force, a strategy that may apply to other receptor systems.

A T cell and APC conjugate creates a similarly constrained organization that drives bending and unbending of the membrane (*35*) with lateral agitation as the TCR snaps open and closed (Fig. 5). Each bend and snap cycle represents a means to initiate T cell activation. Thus, what matters for digital T cell performance isn’t necessarily the presence of a catch bond *per se*, but rather energetically driven signaling through TCR molecular resonance powered by cell surveillance motions and sustained cycles of conformational changes. The catch bond itself may serve more as a gating mechanism to exclude unproductive interactions such as self-reactivity. Bonds that survive this prescreen may then become energized and support molecular resonance with attendant downstream signaling. Local stiffness and other mechanical elements such as the surrounding accessory and adhesion molecules can tune the resonant cycle. Considering that CD8 binds to the side of the MHCα3 domain, the volleying of the TCR will result in a differential yank at the membrane through pMHC-CD8 linkage(s) with potential to repeatedly and simultaneously drive both inside-out and outside-in signaling (*36*).

**Fig. 5.**
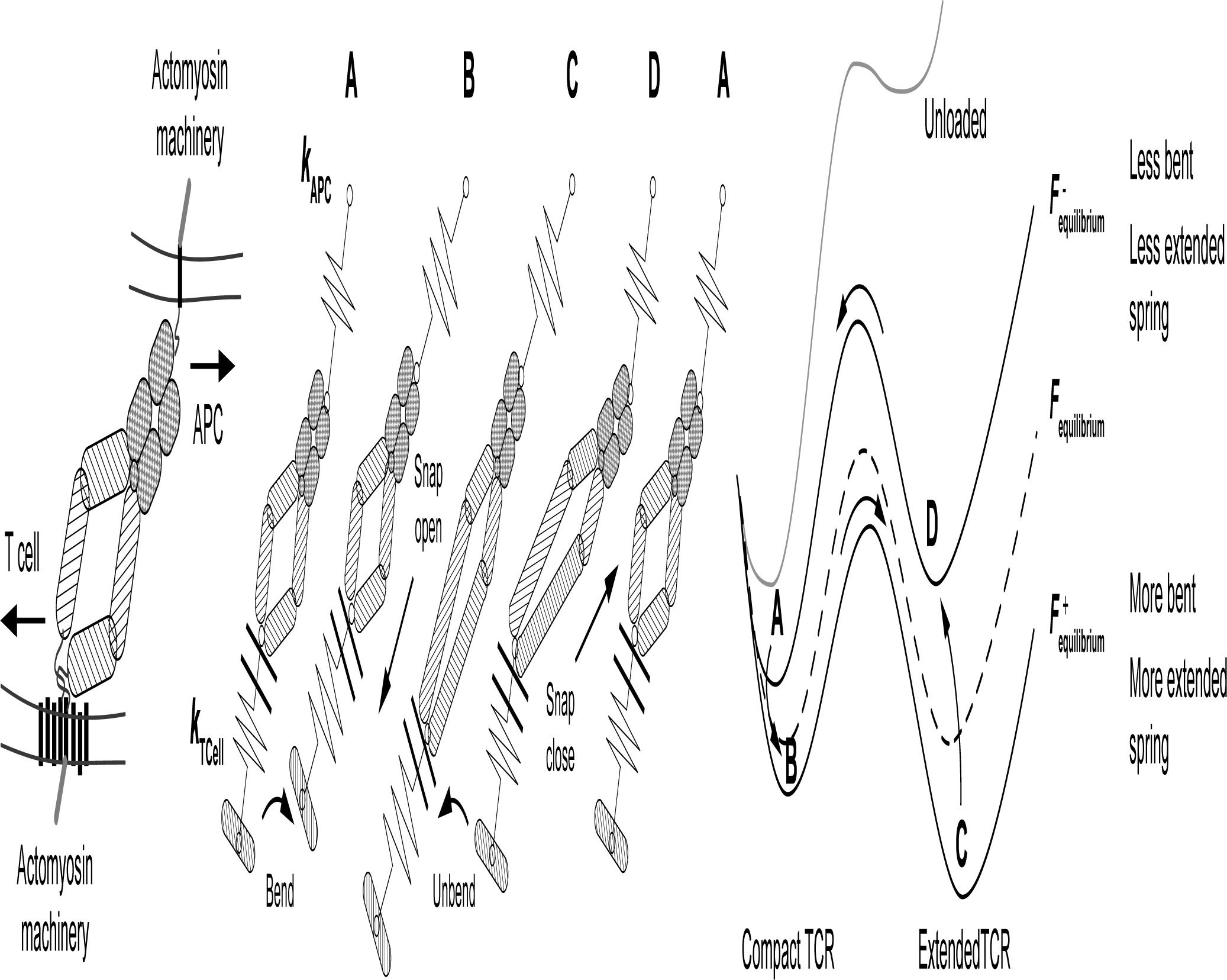
Depiction of the Bend and Snap Cycle (BSC) for sustained signaling associated with TCR molecular resonance. (LEFT) The TCR-pMHC interaction results in formation of a bond between the cell interface with connectivity and force generated through their respective actomyosin machineries. Force across the TCR-pMHC bond is generated through immune surveillance motions arising from T cell pulling to the left, coupling to eight αβTCR membrane associated elements (CD3 ectodomains omitted for clarity) and pMHC coupling to the APC when stationary or pulling to the right. **(MIDDLE)** Various idealized states of the cycle near the equilibrium force are shown with mechanical coupling drawn for simplicity as a “spring” in series with the TCR-pMHC bond. In principle, such mechanical connectivity will have both elastic and viscous character. States B and C depict forces slightly higher than equilibrium while states A and D are slightly lower than equilibrium. States C and D are extended while states A and B are compact. **(RIGHT)** Dynamic energy landscape view of the cycle relative to equilibrium (dashed line) depicting a force slightly higher than equilibrium (F^+^) for states B and C where the system favors transitioning to the extended state. Similarly, states A and D are slightly lower than equilibrium (F^-^) where the closed state is formed. Bend and snap cycle: Force across the bond pulls the system from state A to state B, bending the local membrane shifting the equilibrium to where transitioning to state C is favored. B to C, the TCR snaps open (here shown as an extension of the constant domain) which extends the bond leading to agitation of the membrane. The system immediately adjusts to the new contour length reducing tension on the springs and unbending to state D. The force is now slightly lower than that at equilibrium, reducing tilt of the energy landscape such that the TCR snaps back to state A which retracts the tether agitating the membrane. Cell motions, in turn, pull the bond to state B to allow a repeat of the cycle. Feedback between leftward motions such as the T cell motion indicated (or retrograde flow, not shown) and rightward motion such as from an opposing APC motion (or internal motor activity along the cortical actin, not shown) sustain the cycle. In analogous SM_db_ assays, changes in the contour length due to TCR-pMHC bond extension reduce force across the tether in the optical trap driving the system between a compact higher force state and extended lower force state. Note that without loading of the TCR-pMHC bond (unloaded curve), the energy barrier is too great to overcome. Under load, in contrast, the forward and reverse distance to transition states are relatively balanced which facilitates the bend and snap cycle.

That digital versus analogue performance can be determined by a single amino acid in CDR3 at the αβTCR-pMHC interaction surface (Fig. 1) highlights the impact of the dynamic interactions. Consistent with this notion, single amino acid changes in a peptide (i.e., agonist versus antagonist or null) result in disparate activation of T cells expressing the same TCR and whose TCR-pMHC complexes are virtually identical (*37, 38*). We recently found that asymmetric interdomain-motion and interfacial contacts under load impact the peptide-sensing CDR3 loops, thereby determining mechanical response and peptide discrimination (*39*).

In a digital response, repetitive transitions of single αβTCRs can activate motor proteins to foster kinapse initiation and subsequently mature immunological synapse formation. Both OT and super resolution microscopy experiments revealed that pMHC-ligated αβTCRs and nearby unligated αβTCRs are recruited to initiate TCR clustering (*15, 40*). Analogue TCRs, on the other hand, may benefit from a high density of pMHC ligands to allow for integration of signal or coalescence of smaller TCR clusters as an alternative means to synapse initiation. Regarding TCRs targeting NP_366-374_ /D^b^, both digital and analogue expressing T cells can be engaged productively given the high copy number of that ligand displayed on IAV infected epithelium. Integration of signals from multiple TCRs can afford sensing across a ligand gradient and is presumably operative in T cells bearing either analogue or digital TCRs. By contrast, digital TCRs can recognize sparse ligands like PA_224-233_/D^b^ where analogue TCRs cannot interpret such rare input to drive T cell signaling.

PA TCRs showed distinguishable biophysical performances and activation responses in different functional assays (Figs. 2-3). The PA T-cell transcriptomes were distinct, consistent with the ability of calcium flux and prolonged ERK activation to impact T cell expansion and gene activation (*41, 42*). That PA27 has high levels of the antioxidant Gpx3 and proenkephalin (an attenuator of substance P that promotes asthma via the PI3K/Akt/Nfκb pathway in bronchial epithelium) (*43*) speaks to a potential protective effect mediated by this T cell in lung. The distinct bond lifetime occurring at higher force for PA59 relative to PA27 and PA25 is noteworthy since local tissue stiffness varies 100-fold in normal versus inflammatory conditions, and even more profoundly in different tissue types (*44–46*). The increased breadth of the PA59 catch bond curve may capitalize on accessing higher force interactions expected in stiffer tissues where a bond captured at ∼20 pN can persist long enough to subsequently relax to a resonant state. Perhaps PA59 functions best in stiff locales such as intraepithelial sites where certain resident memory T cells (T_RM_) reside. That mLN PA59 T cells express higher *Itgae* (CD103) and *Itgb7* transcripts than PA27 (data S4), may enhance αEβ7 integrin expression to facilitate intraepithelial site localization through E-cadherin counter-receptors (*47*). Although there is not a universal bioforce load for all αβTCRs (Fig. 2B), the PA25 performance is least ideal among the three PA TCRs examined. Nevertheless, *in vivo* studies have revealed that PA25 function in the lung is broadly comparable to that of the other TCRs, emphasizing how acute inflammation-related cuing fosters productive responses to benefit the host.

Our discovery of digital-vs-analogue performances coupled with distinct bioforce profiles has broad implications. We have identified various quantitative TCR performance metrics including the parameterization of catch bond curves, conformational transitions, critical force and lifetime of volleying, tether formation, activation threshold, and signaling profiles. Additionally, our findings suggest opportunities for nuanced adoptive T-cell immunotherapies and/or cancer vaccine elicitation of relevant TCR specificities with digital TCR performance requirements mandated by sparse neoantigen arrays on cancer cells. The stiffness of desmoplastic solid tumors such as pancreatic cancers likely necessitates utilizing TCRs with greater force-bond lifetime maxima than would compliant hematopoietic tumors.

Biophysical parameterization and functional activation assays in tandem suggest that it is possible to uncover biomarkers of significance. Of note, surface CD3 loss and CD69 upregulation observed here were shown to be associated with TCR quality in a recent independent study (*48*). Our transcriptomic analysis advocates that cytokine receptors and elements involving the cytotoxicity machinery, ribosomes, RNA regulation, and TCR signaling (Fig. 4E) might all be facile biomarker candidates. Defining further the structural basis of TCR performance is worthy of future investigation. Utilizing molecular dynamics in conjunction with machine learning could achieve an aspirational goal of predicting each TCR’s performance within a T-cell repertoire specific for a given pMHC under various physical loads based on primary sequence information.

## Acknowledgements

We gratefully acknowledge Dr. Pasi A. Jänne for facilitating real-time killing assay and Dibyendu K. Das for initial single molecule analysis. We thank Dr. Bruce B. Reinhold for insightful discussion and Ms. Erin C. Mancini and Ms. Jenna K. Koenig for technical assistance and mouse maintenance. All monomers and the tetramers listed in data S5 were obtained through the NIH Tetramer Core Facility.

## Author contributions

Conceptualization: AA, EK, KNB, RJM., JSD-C, WH, MJL, ELR

Methodology: AA, EK, YF, HMS, KNB, DJM, EHK, RJM, JSD-C, WWF, ELH, MJL, ELR

Performing experiments: AA, EK, YF, HMS, KNB, DJM, EHK, RJM, JSD-C, VC, WWF, ELH, JJL, KJZ

Data analysis: AA, EK, YF, HMS, KNB, DJM, EHK, MAB, RJM, JSD-C, VC, WWF, ELH, JJL, KJZ, MJL, ELR

Mouse maintenance: AA, VC

Writing-original Draft: AA, MJL, ELR

Writing-review & editing: AA, EK, YF, HMS, KNB, DJM, EHK, JSD-C, RJM, ELH, WH, MJL, ELR

Funding Acquisition: MJL, ELR Supervision: MYT, MJL, ELR

## Data and materials availability

All sequence files have been deposited in NCBI Gene Expression Omnibus (GEO) under accession GSE240414. Correspondence and requests for materials should be addressed to

All data are available in the main text or the supplementary materials.

