## Supplementary Materials for "Parsing digital or analogue TCR performance through piconewton forces"

**The PDF file includes:**

Materials and Methods
Figs. S1 to S10
References

**Other Supplementary Materials for this manuscript include the following:**

Movies S1 to S3
Data S1 to S7

**Materials and Methods**

Mice and IAV infection

C57BL/6N (B6), B6.129S6-Rag2$^{tm1Fwa}$N12 (*Rag2*$^{-/-}$), and B6.SJL-*Ptprc*$^{a}$/BoyAiTac (CD45.1) mice were purchased from Taconic Biosciences, Inc., housed and bred under specific pathogen-free condition at the DFCI Animal Facility, accredited by the Association for Assessment and Accreditation of Laboratory Animal Care (AAALAC). Euthanasia was performed by $CO_2$ inhalation followed by cervical dislocation. Sex-matched mice were used for each experiment. No gender preference was expressed for this study, and the gender in each experiment was not deliberately selected. Mice at 6-10 weeks of age were infected intranasally with $3x10^4$ $EID_{50}$ of Influenza A/PR/8/34 virus (PR8, H1N1, Charles River Laboratories) as a primary infection under anesthesia with intraperitoneally injection of ketamine/xylazine (120 mg/kg ketamine, 10 mg/kg xylazine) PR8-infected mice were rechallenged intranasally with $5x10^7$ $EID_{50}$ of serologically distinct X:31, A/Aichi/68 (X31, H3N2, Charles River Laboratories). For detecting cell proliferation *in vivo*, mice were intraperitoneally injected with 0.5 mg EdU (Invitrogen) 3 hours before sacrificed. All mouse maintenance, breeding, and experimental procedures were approved under Dana-Farber Cancer Institute Institutional Animal Care and Use Committee (IACUC) protocols 04-113.

Cell lines and cell culture

BW5147.3 cells and CD3δγεζ pMIY vector were a gift from the Vignali lab (St. Jude Children's Research Hospital, Memphis, Tennessee), and mCD8αβ$^{+}$ BW5147.3 cells were generated as previously described(*20*). The cells were maintained in D10 (DMEM medium (Gibco), 10% fetal bovine serum (FBS) (Sigma-Aldrich), 100 IU Penicillin and 100μg/mL

Streptomycin (Corning), 2 mM L-glutamine (Corning) and 55 μM 2-mercaptoethanol (Gibco)), with 400 μg/mL Hygromycin B (Gibco) and 400 μg/ml Geneticin (Gibco). R8 cells were a gift from the Glimcher lab (Harvard School of Public Health, Boston, Massachusetts) and maintained in R10 (RPMI-1640 (Gibco), 10% FBS, 100 IU Penicillin and 100 μg/mL Streptomycin, and 55 μM 2-mercaptoethanol). The retroviral packaging cell line, Plat-E was purchased from Cell Biolabs and were grown in D10 with 1μg/ml puromycin (Sigma-Aldrich) and 10 μg/mL blasticidin (Gibco). LET1 cells were obtained from BEI Resources and maintained in D10. All cells were grown in a 5% $CO_2$ incubator at 37°C.

Cell isolation

Resident $CD8^+$ T cells from lung were isolated as previously described (*49*). Briefly, mice were intravenously injected with 0.8 μg PE-Cy7-conjugated anti-CD8α mAb in 200 μL PBS 5 minutes before euthanized to distinguish $CD8^+$ T cells residing in lung tissue from those in lung vasculature (*50*). Subsequently, lung blood vessels were gently perfused with 60 mL PBS through the right ventricle to wash out the residual injected antibody and then lung tissues were harvested. After mincing lungs with scissors, the chopped tissues were digested with 2 mg/mL collagenase D and 80 U/mL DNaseI in HBSS at 37°C for 1 hour with manual rotation every 10 minutes. Digested tissues were dissociated by gentleMACS Dissociator (Miltenyi Biotec), and the cells were filtered through a 70 μm cell strainer. Red blood cells were eliminated from the cell suspension by treating with hemolysis buffer (140mM $NH_4Cl$, 17mM Tris-HCl, pH7.2). Cell suspensions were resuspended with FACS buffer (2% FBS/0.05% $NaN_3$/PBS) for the following experiments. Mediastinal lymph node (mLN) was harvested, mashed with 3ml syringe on 24-well plate with

RPMI-1640, filtered with 80 μm mesh, washed with RPMI-1640, and resuspended with FACS buffer.

Flow cytometry and cell sorting

Cell suspensions from tissues were first treated with anti-mouse CD16/CD32 mAbs in FACS buffer to block FcR binding for 10 minutes at 4°C and then stained with antibodies indicated in each figure legend in FACS buffer for 20 minutes at 4°C. All antibodies and the concentration used are listed in data S5. For single-cell RT-PCR, the cell suspension from pooled lungs from 2 mice were stained with APC-conjugated $PA_{224-233}$ /H-$2D^b$ tetramer (MBL International Corporation) and PE-conjugated $NP_{366-374}$ /H-$2D^b$ tetramer (MBL International Corporation) for 30 min at RT followed by staining with CD8β mAb to discriminate tissue-resident CD8 T cells defined as $CD8\alpha^-CD8\beta^+$ cells which were spared from vascular $CD8\alpha^+CD8\beta^+$ cells. $PA_{224-233}$- and $NP_{366-374}$-tetramer$^+$ cells were concurrently sorted after gating on 7-Aminoactinomycin $D^-CD8\alpha^-$ $CD8\beta^+$ cells. For RNA-Seq, mCherry$^+$ (PA25), BFP$^+$ (PA27), and GFP$^+$ (PA59) cells were simultaneously sorted after gating on Zombie-NIR$^-$ $CD8\alpha^-CD8\beta^+$ cells.

EdU staining was performed using Click-iT Plus EdU Alexa Fluor 647 or Pacific Blue Flow Cytometry Assay Kit (Invitrogen) following the manufacturer's instructions. For intracellular Ki67 staining, cells were fixed with 4% PFA/PBS, treated with a permeabilization buffer (0.1% saponin (Sigma-Aldrich) in FACS buffer), and incubated with APC-anti mouse-Ki67 mAb at 4 °C for 30 minutes. For detecting NP protein in LET1 cells, the cells were infected with PR8 in Opti-MEM (Gibco) for 1 hour, and then the inoculum was washed out with washing buffer (5 mM $Cacl_2$/5 mM $MgCl_2$/20 mM HEPES/HBSS) followed by culture in R10 for 24 hours. Intracellular NP staining was performed using BD Cytofix/Cytoperm™ Fixation/Permeabilization Kit (BD

Biosciences) and FITC-conjugated Influenza A NP mAb (Invitrogen). Cells were analyzed on a BD LSRFortessa™ Cell Analyzer (BD Biosciences) or sorted by using FACS Aria II (BD Biosciences). Data were analyzed with FlowJo software (FlowJo, LLC).

<u>Single-cell RT-PCR</u>

Single $PA_{224-233}$- or $NP_{366-374}$-specific cells were sorted to each well of 96-well plate that contained 3 μL RT-mix1 (data S6). Amplification of cDNA of TCRα or β was performed using single-cell RT-PCR method as previously described (*51, 52*). All the PCR primers and the component of all the reaction mixtures are listed in data S6. To perform the RT reaction, 2 μL of the RT-mix2 was added to each well containing a single T-cell. After incubation for 60 min at 42 °C, 15 μL of the $1^{st}$ PCR-mix was added to each well to perform the $1^{st}$ PCR reaction. The program for the $1^{st}$ PCR reaction was as follows: 1 min at 98 °C followed by 30 cycles of 10 sec at 98 °C, 5 sec at 53 °C and 40 sec at 72 °C. The resultant $1^{st}$ PCR products were diluted 10-fold with nuclease-free water (Invitrogen) and used for a second cycle of PCR. In the second cycle, TCRα and β were amplified separately. To amplify the cDNA of TCRα or TCRβ, 2 μL of the diluted $1^{st}$ PCR products was added to each well of new 96-well PCR plate containing 18 μL of the $2^{nd}$ PCRα-mix or the $2^{nd}$ PCRβ-mix, respectively. The program for the $2^{nd}$ PCR reaction was as follows: 1 min at 98 °C followed by 35 cycles of 10 sec at 98 °C, 5 sec at 58 °C and 30 sec at 72 °C. The $2^{nd}$ PCR products were then analyzed with the Ca_RV3 primer for TCRα or Cb_RV3 primer for TCRβ by direct sequencing. The TCR repertoire was analyzed with reference to the IMGT (http://www.imgt.org).

Clonotype definition

Assembled TCR genetic elements yielded nucleotide sequences encoding a subunit clonotype for each cell. TCRα and TCRβ clonotypes were counted separately and those clonotypes with productive CDR3s were selected for analysis. Each paired TCRαβ clonotype unique to a given single cell was of particular functional interest. The hierarchy of subunit clonotypes is shown in the pie charts in Fig. 1A and Fig. 2A. TRV and TRJ repertoires and CDR3 sequence for all TCR clonotypes are listed in data S1.

Retrovirus production and transduction

cDNA encoding TCRβ-P2A-TCRα was inserted into a retroviral vector pMSCV-IRES-GFP II (pMIG II, Addgene, #52107), pMSCV-IRES-mCherry FP (Addgene, #52114), or pMSCV-IRES-Blue FP (Addgene, #52115). For the real-time killing assay, non-fluorescence TCRαβ-expressing vectors were made by cutting IRES-EGFP site out from pMIG II with restriction enzymes and by inserting TCR cDNA. The vector was transfected into Plat-E cells with Fugene HD (Promega). Retrovirus in the cultured supernatant was collected 72 hours later and frozen at -80°C until use. Thawed retrovirus was transduced with Retronectin (Takara Bio) using Retronectin-bound virus infection methods according to the manufacturer's instruction. Retrovirus was transduced into mCD8αβ$^{+}$ BW5147.3 cells to generate TCRαβ-expressing cell lines or into mouse bone marrow (BM) cells to generate Rg mice. To establish TCRαβ-expressing BW cell lines, the transduced cells were sorted by FACS Aria II (BD Biosciences) to match the TCR expression based on the cell surface expression of CD3ε. Before all assays were performed, CD3 or TCRβ surface level was confirmed to be matched in a group.

Generation of Rg mice

Rg mice were generated as previously described (*53*). Briefly, BM cells were harvested from *Rag2*$^{-/-}$ mice, and hematopoietic stem cells (HSC) were enriched by EasySep Mouse Hematopoietic Progenitor Cell Isolation Kit (STEMCELL Technologies) followed by expansion in Stem cell medium (StemPro™-34 SFM (Gibco), 5% FBS, 100 IU Penicillin and 100 mg/mL Streptomycin, 2 mM L-glutamine, with 50 ng/mL mIL-3 (STEMCELL Technologies), 50 ng/mL hIL-3 (STEMCELL Technologies) and 50 ng/mL mSCF (STEMCELL Technologies)) for 3 days. Subsequently, HSC were transduced with retrovirus encoding TCRαβ, cultured in Stem cell medium for 3 days, and then transferred into *Rag2*$^{-/-}$ mice irradiated with Gamma Cell 40 $Cs^{137}$ Irradiator (Thratonics) one day before. Rg mouse blood was analyzed for CD8 development by flow cytometry 6 weeks after transplantation, and the mice expressing more than 10% of CD8 T cells in $CD45^+$ cells were used for generation of RgC mice.

Generation of RgC mice

For single RgC mice, peripheral LNs and spleen were harvested from Rg mice, and $FP^+CD8\beta^+CD44^-$ naïve T cells were sorted. Subsequently, $2\sim10\times10^4$ cells were intravenously transferred into recipient B6 mice. For mixed RgC mice, an equal number of Rg T cells were mixed before transfer. The ratio of the mixed Rg T cells was confirmed by the combination of FP and Vβ expression by flow cytometry. RgC mice were intranasally infected with PR8 one day after adoptive transfer. For real-time killing assay, $CD8\beta^+CD44^-$ naïve T cells from non-FP Rg mice were sorted and adoptively transferred into recipient CD45.1 mice.

TCR protein expression

$PA_{224}$-specific TCRαβ or $NP_{366}$-specific NP41αβ Leucine Zipper (LZ) proteins were produced and used for SM assay as previously described (*17, 54*), Briefly, separate chains were expressed in Expi293F (ThermoFisher) cells according to the manufacturer protocol and purified from supernatants as a LZ paired heterodimer utilizing an anti-LZ mAb (clone 2H11). The TCRαβ constructs consist of V and C ectodomains connected to the 30-amino acid LZ motif via a 15-residue flexible linker sequence. The heterodimer was covalently linked via the native disulfides located at the C-terminal end of each ectodomain.

SMSC assay

SMSC assay was performed to measure the specific bond lifetime of TCR-pMHC interaction using a single-molecule DNA tether, which was functionalized with a half anti-biotin antibody to capture biotinylated mutated-pMHC at one end and with a digoxigenin tag for tether adhesion on anti-digoxygenin-coated polystyrene beads (1.0 µm in diameter, Spherotech Inc.) at the other end. The bead slurry was washed with PBST buffer (1X PBS + 0.02% (v/v) Tween-20) twice and then resuspended with a dilution factor of 200X in colorless DMEM medium supplemented with 5 mg/mL bovine serum albumin (BSA) for bond lifetime measurements. Cells used in SMSC assay were rinsed once with colorless 1 mL of DMEM medium and re-suspended to a final concentration of $2x10^6$ cells/mL. Next, 20 µL of this cell suspension was transferred into the flow chamber, where the cells were allowed to attach to a polylysine coated coverslip. The chamber was then incubated at 37 °C with 5% $CO_2$ for 30 minutes. Afterward, the coverslip surface was passivated using colorless DMEM medium supplemented with 5 mg/mL BSA, followed by a 10-minute incubation at 37°C with 5% $CO_2$. Subsequently, approximately 20 µL of pMHC-tethered bead slurry was pipetted on one side of the flow chamber and sucked out the other side by capillary action using a Kimwipe. The pMHC-tethered bead was trapped by the trapping laser (1064 nm) and brought close to a nearby cell, resulting in the formation of a stable tether between the bead and T cell. Pulling force was generated by stepping the piezo stage with a defined distance in the direction opposite to the approaching direction. Specific procedures used for preparing the beads and measuring bond lifetime are described in a previous work (*17*).

SCAR assay

SCAR assay was performed to measure early T-cell activation via staining the cells with Quest Rhod-4, AM (AAT Bioquest, Inc.) to visualize the intracellular $Ca^{2+}$ flux. Subsequently, WT pMHC-coated beads with varying interfacial copy numbers were employed to assess the triggering capability for different T cell lines. Compared to the normal forces in SMSC assay, tangential forces parallel to the cell-bead interface were applied with defined magnitude. Detailed protocols for the preparation and quantification of interfacial number of molecules of pMHC-coated beads, Quest Rhod-4 staining, and intracellular $Ca^{2+}$ activation with optically trapped beads can be found in a previously published work (*15*). For Fig. 1D and fig. S2F, G, adaptations made to the protocol include the following: streptavidin beads with a diameter of 1.36 μm (Spherotech Inc.) were used. Poly-L-lysine coated coverslips were used to facilitate cell binding to the coverslip surface. Flow channels were formed with a single layer of double-sided tape. Fluorescence images were taken every 5 seconds for 10-15 minutes.

SCAR experiments performed for Fig. 1D and fig. S2F, G were acquired using a microscope adapted for combined trapping and single molecule fluorescence with low levels of fluorescence excitation. In these experiments we used a trapping laser power of ~350 mW and total fluorescence excitation laser power of 5 μW in Epi mode with an excitation zone spread over an area of ~2,827 $\mu m^2$ (~$1.77*10^{-3}$ $\mu W/\mu m^2$).

SM assay

Purified single heterodimers with LZ as described above coated one bead via 2H11, which was covalently linked to 1.23 μm polystyrene beads (Spherotech Inc.) via EDC chemistry. A 3500 bp DNA tether with digoxigenin on one end and streptavidin on the other was used to link biotinylated mutant $PA_{224-233}$/H-$2D^b$ or $NP_{366-374}$/H-$2D^b$ monomer to a second 1.23 μm polystyrene bead coated with anti-digoxigenin. Streptavidin was used to connect limiting amounts of pMHC to the DNA tether. Beads were loaded into separate channels of the microfluidics system on the LUMICKS m-Trap. Each bead was trapped in respective channels and moved to empty PBS channels where they were calibrated, brought together to form tethers, and were rapidly pulled apart to load a known force. Beads were used for 3-7 lifetime measurements before discarding and trapping a new bead pair. Trap stiffnesses ranged from 0.20-0.30 pN/nm. Interactions were measured until bond rupture. In the case of volleying the trap separation could be slightly adjusted by a few nm to cause fraction of a pN changes in force to keep the system within the volleying force range. Changes in positional distribution could be directly observed as a result of said minute adjustments (Fig. 3H).

Tetramer binding assay

$5x10^5$ PA-or NP-TCR transduced BW cells were plated on 96-well plates, and titrated WT- or CD8BS-mutant $PA_{224-233}$/H-$2D^b$, or WT- or mutant $NP_{366-374}$/H-$2D^b$ tetramer in FACS buffer was added, respectively. The cells were incubated at RT for 30 minutes, washed with FACS buffer twice, and the fluorescence intensity was analyzed by flow cytometry. The tetramers used in each experiment are listed in data S5.

Tetramer dissociation assay

5x$10^5$ PA-BW cells were treated with mutant $PA_{224\text{-}233}$/H-2$D^b$ tetramer at RT for 30 minutes. WT-$NP_{366\text{-}374}$/H-2$D^b$ tetramer was used for NP-BW cells due to an inability of the mutant tetramer binding for NP41. The cells were washed and plated on 96-well plates. Subsequently, 3 μg Fab fragment of anti-mouse H-2$D^b$/H-2$L^d$ antibody was added to the cells for the indicated times at RT, and then the cells were immediately fixed in 4% PFA/PBS. The cells were washed, and the fluorescence intensity of the tetramer was analyzed by flow cytometry. The cells without Fab addition were used as a control of time 0. The tetramers used in each experiment are listed in data S5. Fab fragment of anti-mouse H-2$D^b$/H-2$L^d$ mAb (BioLegend) was made by Pierce™ Fab Preparation Kit (Pierce) according to the manufacturer's instruction.

Tetramer activation assay

For pERK assay, the cells were resuspended at a concentration of 1 x $10^6$ cells/mL in a final volume of 0.5 mL. For each sample, either the WT $PA_{224\text{-}233}$/H-2$D^b$ tetramer (0.75 μg/mL) or the WT $NP_{366\text{-}374}$/H-2$D^b$ tetramer (2.75 μg/mL) was added to the PA- or NP-BW cell samples, respectively. Control samples were also prepared for each cell line without the addition of tetramer at time 0. All cell samples were then incubated on ice for 20 minutes and then washed with DMEM to remove excess tetramer. The cells were resuspended at a final concentration of 2 x $10^6$ cells per 0.1 mL and incubated at 37 °C for the indicated 0–10-minute time points. Lysis buffer (final concentration after dilution: 1% Triton X-100, 0.05% SDS, 50 mM Tris pH 7.4, 150 mM NaCl, 2 mM $NaVO_3$, 1 mM NEM, and Roche cOmplete protease cocktail) was added to the cells and the samples were immediately placed on dry ice to await further processing. Cell samples were thawed for 10 minutes on ice and centrifuged at 13K rpm, 4°C for 15 minutes and clarified lysate was transferred to a clean tube. Aliquots were run on 4%-12% Bis-Tris NuPAGE gels, transferred to PVDF membrane for detection with ERK (W15133B clone) and phosphorylated ERK (4B11B69 clone) antibodies, and imaged using the BioRad ChemiDoc imaging system. Band density was measured utilizing the Image Lab software and the level of phosphorylated ERK was normalized to ERK for each sample. This was then further normalized to the median within each experimental set to mitigate artifacts of antibody staining. Technical replicates were generated for each biological replicate by preparing three sets of cells and conducting at least three biological replicates for each activation assay.

For analysis of CD3ε loss and CD69 upregulation after tetramer activation, titrated WT $PA_{224\text{-}233}$/H-$2D^b$ tetramer or WT $NP_{366\text{-}374}$/H-$2D^b$ tetramer was added to $2x10^5$ PA- or NP-BW cells, respectively. The cells were cultured in D10 at 37°C overnight, washed with FACS buffer twice, stained with anti-mouse CD3ε mAb and anti-mouse CD69 mAb, and analyzed by flow cytometry. The gMFI of CD3ε and CD69 without the tetramer was normalized to 100%.

### Cell-based functional avidity assay

$1x10^5$ PA- or NP-BW cells were cultured in D10 with titrated $PA_{224\text{-}233}$ or $NP_{366\text{-}374}$ peptide (from $1x10^{-5}$ to $1x10^4$ ng/mL), respectively, at 37°C for 16-18 hours overnight. $1x10^5$ R8 cells used as APC were treated with Mitomycin C (Sigma-Aldrich) and washed with PBS three times prior to use. IL-2 concentration in culture supernatant was measured by ELISA assay according to the manufacturer's instruction. $EC_{50}$ of peptide response to each BW cells were calculated by Prism 7 (GraphPad Software).

Real-time killing assay

To visualize LET1 cells, mCherry retrovirus derived from pMSCV-IRES-mCherry FP vector (Addgene) was transduced. $2x10^4$ $mCherry^+$ LET1 cells were seeded 18 hours before PR8 infection on 96 well plate (Corning). The cells were washed with washing buffer (5 mM $Cacl_2$/5 mM $MgCl_2$/20mM HEPES/HBSS) and infected with titrated PR8 (from $4x10^5$ $EID_{50}$ to $4x10^8$ $EID_{50}$) in Opti-MEM (Gibco) at 37°C for 1 hour. Rg T cells from mLN or lung were obtained from CD45.1 RgC mice infected with PR8 seven days before and sorted as $CD45.2^+$ $CD8\beta^+$ $CD44^+$ Zombie $Aqua^-$ intravascular stained $CD8\alpha^-$. $2x10^4$ Rg T cells were labeled with 10 μM Cell Proliferation Dye eFluor™ 450 (eBioscience) and plated on the infected $mCherry^+$ LET1 cells. CellEvent Caspase-3/7 Green ReadyProbes™ Reagent (Invitrogen) was added to the wells to measure apoptosis. Plates were housed in a BioSpa 8 Automated Incubator (Agilent) and were scanned at regular intervals using a Cytation 5 Cell Imaging Multimode Reader (Agilent). Image processing and data analysis was conducted using the Gen5 3.11 software (Agilent). Whereas the Cell Proliferation Dye eFluor™ 450 remains bound to apoptotic T cells, mCherry was quickly lost from apoptotic LET1 cells due to intracellular expression of the soluble protein. Therefore, LET1-specific killing was quantified by subtracting T cell-specific apoptotic signal (green and blue overlapping area) from total caspase-3/7 activity levels (total green area). Efficiency of LET1 killing was then calculated via normalization to confluency of viable LET1 cells at each respective timepoint (total red area).

Measurement of virus titer

Viral copy number was determined by the method previously described(*55*). Briefly, lung tissues were collected, preserved in RNAlater (SIGMA), and frozen at -80°C until used. Once thawed, the tissues were homogenized by Tissue disruptor (QIAGEN), and RNA was extracted with PureLink RNA Mini kit (Invitrogen). Reverse transcription was conducted with High-Capacity cDNA Reverse Transcription Kit (Applied Biosystems), and qPCR was performed with PowerUP SYBR Green Master Mix for qPCR (Applied Biosystems) and primers specific for NP (Forward: 5'- GAT TGG TGG AAT TGG ACG: Reverse: 5'- AGA GCA CCA TTC TCT CTA TT-3') using the 7900HT Fast Real Time PCR System (Applied Biosystems). The standard calibration curve for qPCR was obtained by stepwise dilution of the cloned NP gene fragment with a known copy number.

RNA-seq

Following isolation of PA25, PA27, and PA59 Rg T cells from both lung and mLN from 3 mixed RgC mice for each TCR, each preparation was processed individually (i.e. 3 biological replicates) for isolation of total RNA using the RNAqueous-4PCR total RNA isolation kit (ThermoFisher) that minimizes genomic DNA contamination. Cell input for the LN preparations was 11,643 ± 2,486 (mean ± SEM, n =9), and for the lung preparations was 45,327 ± 6,225 (mean ± SEM, n =9). Quality control was assessed using an Agilent 4200 TapeStation. Libraries suitable for RNA-Seq analysis were prepared using the SMART-Seq v4 Ultra low Input RNA kit (Takara), followed by addition of Illumina adapters and 150 bp paired end sequencing on the Illumina NovaSeq 6000 platform (MedGenome Inc.).

RNA-seq data analysis

Output per library for the LN Rg T cells averaged ~60M reads, and for the lung resident Rg T cells averaged ~80M reads. The paired output fasta files for each sample were checked for quality using FastQC (v0.11.8) and adapters trimmed using FastqMcf (v1.05) and Cutadapt (v4.4). Sequenced RNA-Seq reads were aligned to the mouse genome (mm10) using STAR (2.4.2a) (*56*). The RNA-Seq pipeline Viper (*57*) was used to generate gene-level read counts, gene-level TPM (Transcript Per Million) values, and to perform Principal Component Analysis for the analyzed samples. Differentially expressed genes were identified with the R package Deseq2 (1.38.3) (*58*) using fold-change threshold of 2 and adjusted P-value threshold of 0.05. Gene Set Enrichment Analysis (GSEA) was performed using the R package clusterProfiler (4.6.2) (*59*). The sources of gene sets used for the analysis are provided in data S7.

Statistics

Statistical analyses were performed with GraphPad Prism software (v9) except pERK assay and RNA-seq data analysis. Statistical tests used, one-way ANOVA, linear regression, Kolmogorov-Smirnov Test, paired t-test, Wald test, Kruskal-Wallis or unpaired t-test were indicated in Figure legends. For three independent experiments of pERK assay, the statistics were performed by regression using trend line analysis models accounting for interexperimental variability (R Statistical Software Package (v4.1.2).

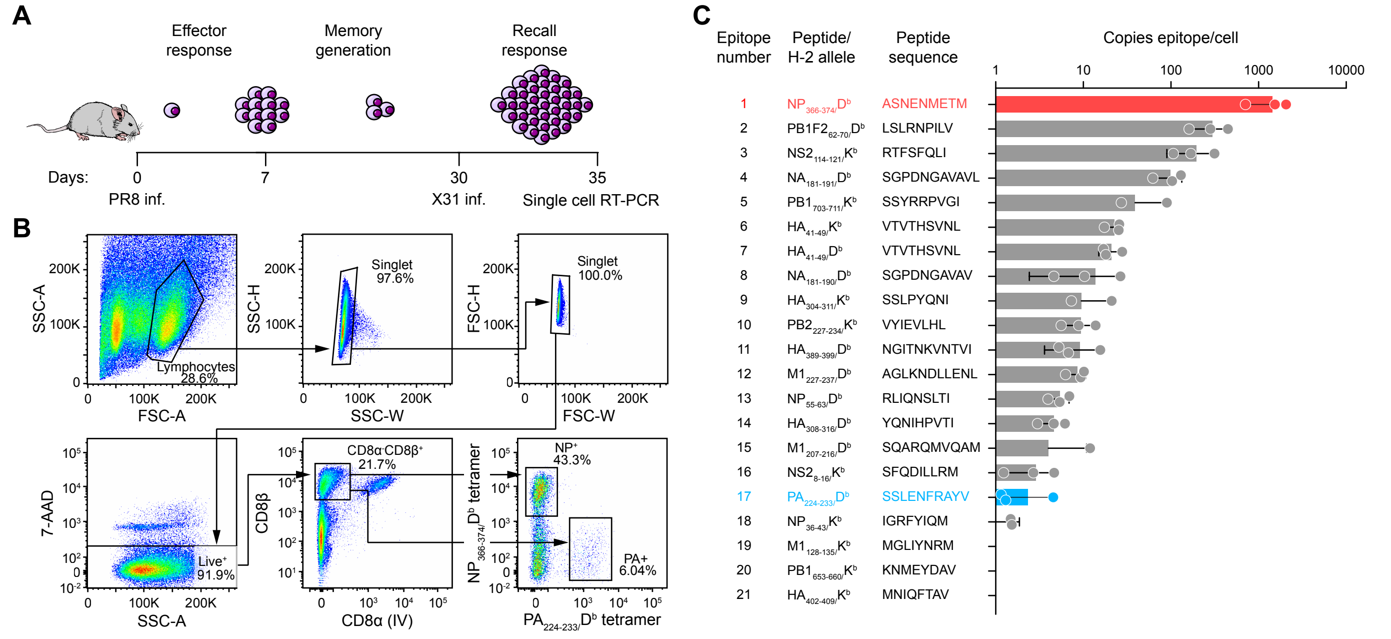

A
Effector response
Memory generation
Recall response
Days: 0 7 30 35
PR8 inf.
X31 inf.
Single cell RT-PCR
B
SSC-A
FSC-A
Lymphocytes 28.6%
SSC-H
SSC-W
Singlet 97.6%
FSC-H
FSC-W
Singlet 100.0%
7-AAD
SSC-A
Live+ 91.9%
CD8β
CD8α (IV)
CD8α- CD8β+ 21.7%
NP$_{366-374}$/D$^b$ tetramer
PA$_{224-233}$/D$^b$ tetramer
NP+ 43.3%
PA+ 6.04%
C
Epitope number
Peptide/ H-2 allele
Peptide sequence
Copies epitope/cell
1 10 100 1000 10000
1 NP$_{366-374}$/D$^b$ ASNENMETM
2 PB1F2$_{62-70}$/D$^b$ LSLRNPILV
3 NS2$_{114-121}$/K$^b$ RTFSFQLI
4 NA$_{181-191}$/D$^b$ SGPDNGAVAVL
5 PB1$_{703-711}$/K$^b$ SSYRRPVGI
6 HA$_{41-49}$/K$^b$ VTVTHSVNL
7 HA$_{41-49}$/D$^b$ VTVTHSVNL
8 NA$_{181-190}$/D$^b$ SGPDNGAVAV
9 HA$_{304-311}$/K$^b$ SSLPYQNI
10 PB2$_{227-234}$/K$^b$ VYIEVLHL
11 HA$_{389-399}$/D$^b$ NGITNKVNTVI
12 M1$_{227-237}$/D$^b$ AGLKNDLLENL
13 NP$_{55-63}$/D$^b$ RLIQNSLTI
14 HA$_{308-316}$/D$^b$ YQNIHPVTI
15 M1$_{207-216}$/D$^b$ SQARQMVQAM
16 NS2$_{8-16}$/K$^b$ SFQDILLRM
17 PA$_{224-233}$/D$^b$ SSLENFRAYV
18 NP$_{36-43}$/K$^b$ IGRFYIQM
19 M1$_{128-135}$/K$^b$ MGLIYNRM
20 PB1$_{653-660}$/K$^b$ KNMEYDAV
21 HA$_{402-409}$/K$^b$ MNIQFTAV

**Fig. S1. Schematic of cell isolation for single-cell RT-PCR and analysis of $NP_{366\text{-}374}/D^b$- and $PA_{224\text{-}233}/D^b$- specific TCR directed at distinct peptides with divergent copy numbers after IAV infection.**

**(A)** Workflow to clone $NP_{366\text{-}374}/D^b$- and $PA_{224\text{-}233}/D^b$- specific TCRs derived from lung resident $CD8^+$ T cells. T cells were isolated 5 days post-recall by cell sorting and single cell RT-PCR. (**B**) Sorting strategy to isolate $NP_{366\text{-}374}/D^b$- and $PA_{224\text{-}233}/D^b$- specific T cells. **(C)** Copy number of 21 peptide/H-2 MHC complexes on LET1 cells infected with Influenza A/PR/8/34 virus (PR8) based upon data adapted from Wu et al. *Nature communications* (2019). The pMHCs were isolated from infected LET1 cells, and then the peptides were eluted from their immunoaffinity-purified $K^b$ and $D^b$ MHCI molecules and analyzed by LC-MS as described. Results from LET1 are summarized, with $NP_{366\text{-}374}/D^b$- and $PA_{224\text{-}233}/D^b$ peptide copy numbers adapted from that publication.

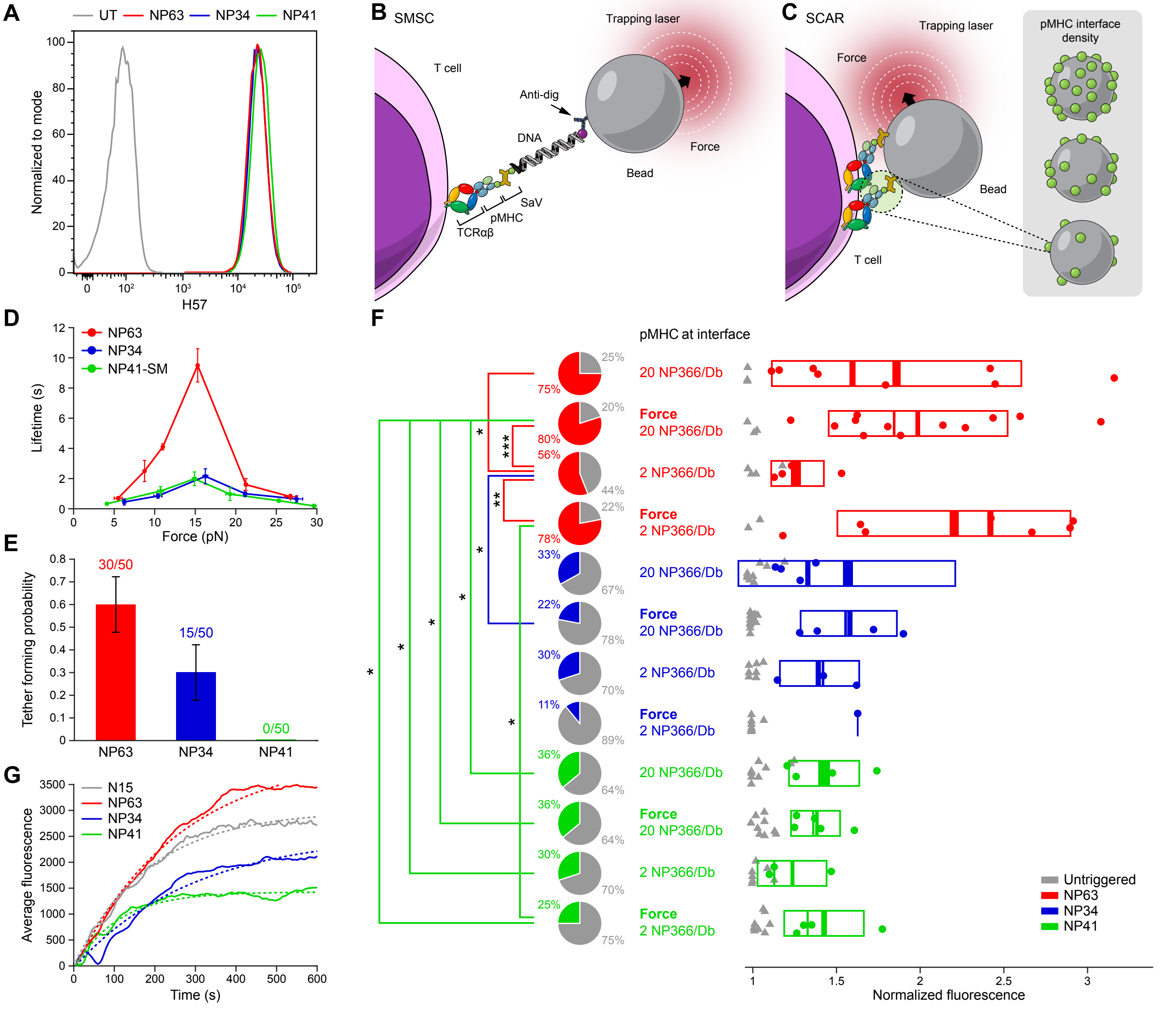

A
UT
NP63
NP34
NP41
Normalized to mode
H57
B
SMSC
T cell
Trapping laser
Anti-dig
DNA
Force
Bead
SaV
pMHC
TCRαβ
C
SCAR
Trapping laser
Force
Bead
T cell
pMHC interface density
D
NP63
NP34
NP41-SM
Lifetime (s)
Force (pN)
E
30/50
15/50
0/50
Tether forming probability
NP63
NP34
NP41
F
pMHC at interface
20 NP366/Db
Force 20 NP366/Db
2 NP366/Db
Force 2 NP366/Db
Untriggered
NP63
NP34
NP41
Normalized fluorescence
G
N15
NP63
NP34
NP41
Average fluorescence
Time (s)

**Fig. S2. The greater activation under force of the digital NP63 TCR relative to analogue NP34 and NP41.**

**A**, TCRβ expression on CD8αβ$^{+}$ TCR$^{-}$ BW5147 cell line transduced with NP34, NP41, and NP63 TCRαβ after cell sorting to match TCR expression as defined by H57 anti-Cβ mAb. (**B**) Single molecule single cell (SMSC) assay design for optical tweezer experiments. Beads functionalized with DNA tethers terminating in pMHC are actively introduced to the cell surface to initiate bond-formation, then retracted for bond-loading. Movement of the bead relative to the cell is controlled by a piezo stage that translates the cell relative to a stationary trapping laser. (**C**) Single cell activation requirement (SCAR) assay design for optical tweezer experiments. T cells are introduced to pMHC coated beads, with varying densities of an agonist pMHC ($NP_{366}/D^b$ or $PA_{224-233}$ $/D^b$), to promote bond formation at the interface. Moving the piezo stage relative to the T cell applies a vectorial force to the system via the optical trap. (**D**) Force vs. lifetime distributions, comparing NP63 and NP34 obtained with SMSC assay to NP41 obtained with SM assay. (**E**) Tether forming probability in the single-molecule single-cell (SMSC) assay under identical bead and coverslip conditions for each clone. At conditions where sufficient tethers are formed between $NP_{366}/D^b$ pMHC and NP63 or NP34 cells, no tethers are formed in the case of NP41. (**F**) SCAR assay used to measure the calcium flux in NP63-, NP34-, and NP41-BW cells using either 2 or 20 interfacial copy number of $NP_{366}/D^b$, conducted on a high-resolution dual fluorescence-OT microscope (see SCAR methods for more details). Experiments were conducted with and without external force application as indicated. The calcium flux is represented as ratio between the maximum fluorescence intensity ($I_{max}$) and the initial fluorescence intensity ($I_0$). Grey triangles represent cells that did not trigger after bead introduction, while filled circles represent cells that did trigger. Some cells are designated as non-triggered cell even though their values are higher than a triggered cell due to the shape of the calcium flux profile. The profile of those non-triggered cells rises and then falls back below baseline in quick succession (within ~1-2 minutes), whereas the lower values considered “triggered” have sustained calcium flux profiles, and thus extended signaling. The width of the rectangles shows the SD with the mean and median as the thick and thin line, respectively. Pie charts show triggering percentage of each population, where colored wedges are the triggering cells. One-way ANOVA used to quantify significant difference in triggering normalized fluorescence means; ***$P<0.001$, **$P<0.01$, *$P<0.05$. **(G)** Average fluorescence curves of SCAR data from the high-resolution microscope with the baseline fluorescence subtracted out (solid line). Individual curves from all triggering cells from each respective cell line were pooled (combining interfacial pMHC concentrations of 20 and 2 with force for NP63, NP34 and NP41 and 2 with force for N15) and averaged at time zero. All curves fit to $y = A * \left(1 - e^{-x/t}\right)$, where y is the fluorescence intensity, A is the amplitude, *t* is the rise-time constant (s), and x is the time (s) (dashed line). The amplitudes and 95% confidence intervals for NP63, NP34, NP41, and N15 are 4,085 ± 106, 2,547 ± 146, 1,425± 22, and 2,993± 47, respectively. The rise-time constants and 95% confidence intervals for NP63, NP34, NP41, and N15 are 261 ± 14.8, 293 ± 34.1, 108 ± 6.9, and 185 ± 8.1 seconds, respectively. N15 recognizes VSV8/$K^b$, an epitope derived from vesicular stomatitis virus[38], unlike the NP TCRs which bind $NP_{366}/D^b$.

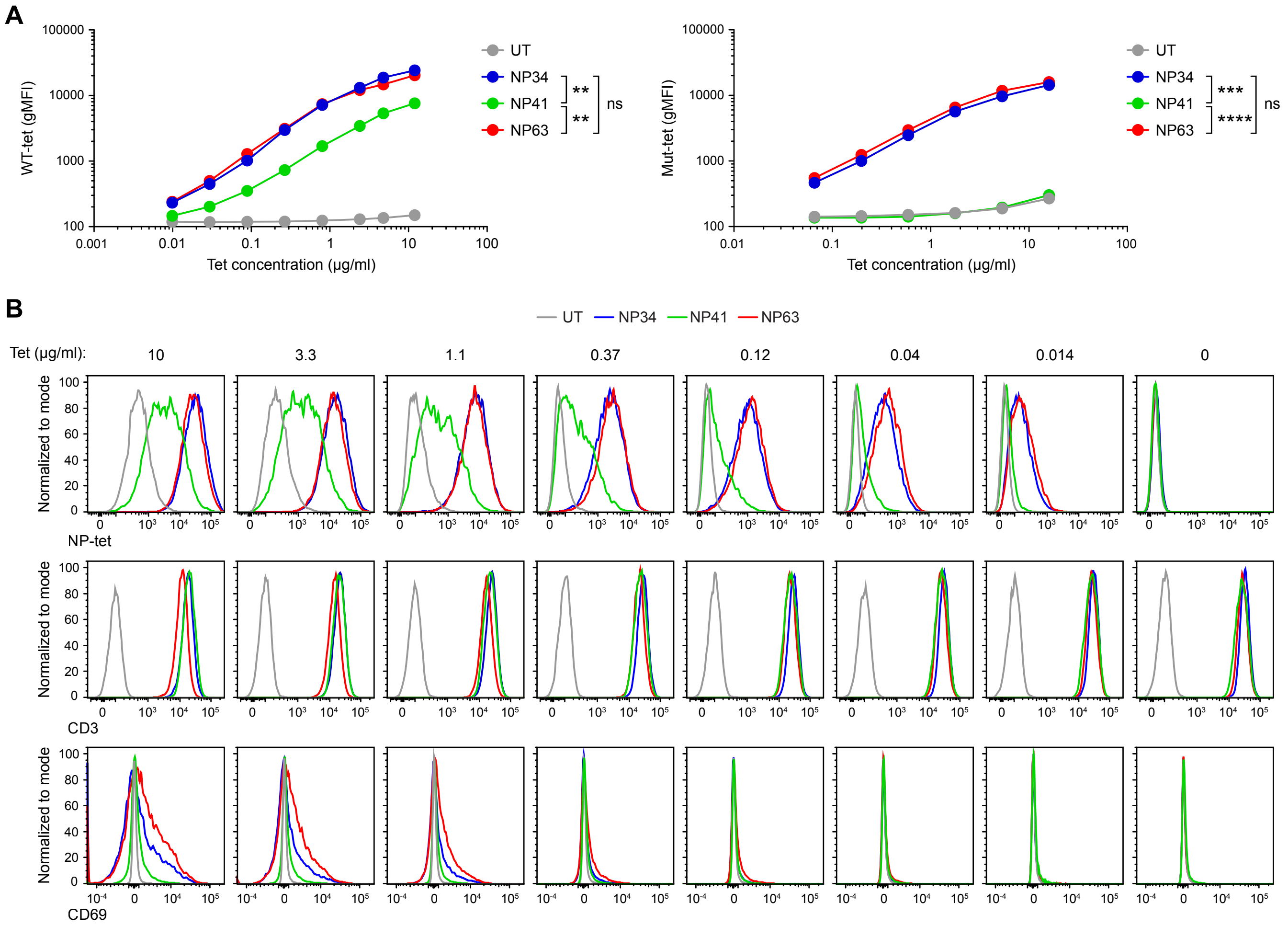
A
WT-tet (gMFI)
Tet concentration (µg/ml)
UT
NP34
NP41
NP63
**
**
ns
Mut-tet (gMFI)
***
****
B
Tet (µg/ml):
10
3.3
1.1
0.37
0.12
0.04
0.014
0
Normalized to mode
NP-tet
CD3
CD69

**Fig. S3. Superior activation ability of T cells expressing digital NP63 TCR in response to tetramer stimulation *in vitro*.**

**(A)** Tetramer binding of NP$_{366\text{-}374}$ /D$^b$ WT-tetramer (left) and CD8BS-mutant tetramer (right) to NP34-, 41-, and 63-BW cell lines and BW UT, cells untransduced with a NP-specific TCR. Tetramer treatment was performed at 20 °C for 30 minutes. Data are representative of two independent experiments. ****P <0.0001, **P<0.01; ns, not significant. P values were calculated by linear regression. **(B)** Histogram of tetramer (top), anti-CD3 (middle), and anti-CD69 binding (bottom) for NP34 (blue)-, 41 (green)-, and 63 (red)-BW cell lines after stimulation with indicated concentration of NP$_{366\text{-}374}$ /D$^b$ WT-tetramer at 37 °C overnight, as shown in Fig. 1F, I and J, respectively. Data are representative of three replicates in two independent experiments.

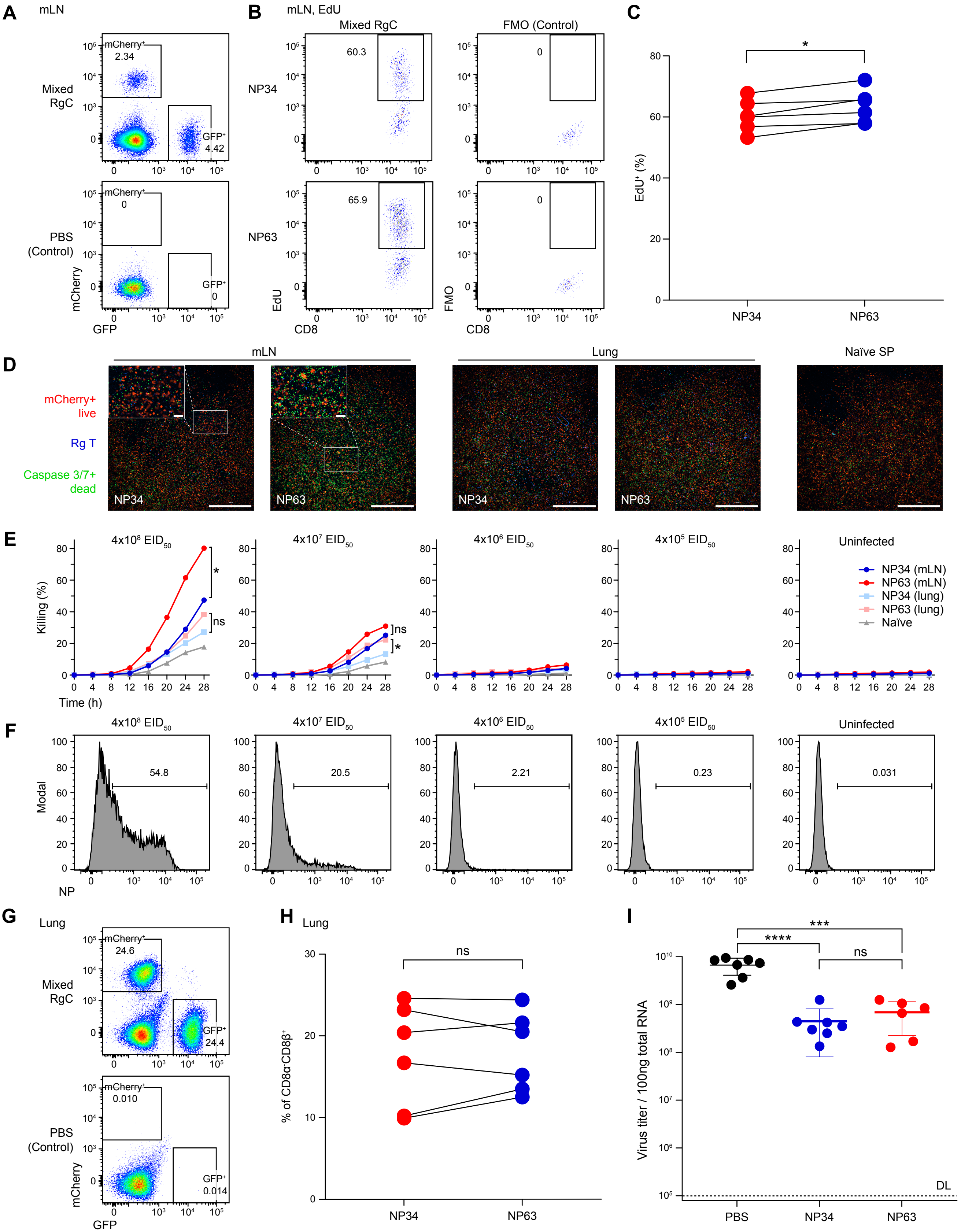
A
mLN
Mixed RgC
PBS (Control)
mCherry+
2.34
GFP+
4.42
mCherry+
0
GFP+
0
mCherry
GFP
B
mLN, EdU
Mixed RgC
FMO (Control)
NP34
NP63
60.3
65.9
0
0
EdU
FMO
CD8
C
*
EdU+ (%)
NP34
NP63
D
mLN
Lung
Naïve SP
mCherry+ live
Rg T
Caspase 3/7+ dead
NP34
NP63
NP34
NP63
E
4x10^8 EID50
4x10^7 EID50
4x10^6 EID50
4x10^5 EID50
Uninfected
Killing (%)
Time (h)
*
ns
ns
*
NP34 (mLN)
NP63 (mLN)
NP34 (lung)
NP63 (lung)
Naïve
F
4x10^8 EID50
4x10^7 EID50
4x10^6 EID50
4x10^5 EID50
Uninfected
Modal
NP
54.8
20.5
2.21
0.23
0.031
G
Lung
Mixed RgC
PBS (Control)
mCherry+
24.6
GFP+
24.4
mCherry+
0.010
GFP+
0.014
mCherry
GFP
H
Lung
ns
% of CD8α+CD8β+
NP34
NP63
I
***
****
ns
Virus titer / 100ng total RNA
DL
PBS
NP34
NP63

**Fig. S4. NP34 and NP63 Rg T cell activation *in vivo* after IAV infection.**

**(A)** Representative FACS plots of NP34 ($mCherry^+$) and NP63 ($GFP^+$) T cells in mLN of mixed RgC mice 7 days after PR8 infection shown in Fig. 1K. Data were derived after first gating on $CD8b^+$ cells. Control mice were injected with PBS without Rg T cells when the RgC mice were generated. (**B, C)** EdU incorporation in NP34 and NP63 T cells. Representative FACS plots (**B**) and quantification (**C**) are shown. (**D**) Original images of mLN Rg T cell- (top) and lung Rg T cell (bottom)-mediated killing of LET1 cells infected with $4x10^8$ $EID_{50}$ PR8 at 20 hours, as shown in Fig. 1M. Naïve splenic T cells were used as control (Naïve SP) of uninfected B6 mice (bottom right). Rg T cells were sorted from six pooled RgC mice that received a single RgT cell type (dpi 7) and then cultured on infected LET1 cells. LET1 cells are visualized by transduced mCherry, apoptotic cells are visualized in green using Caspase-3/7 Green ReadyProbes, and Rg T cells are stained in blue with Cell Proliferation Dye eFluor 450. mLN NP63 Rg T cells show a significant killing ability indicated by increased green apoptotic signals and loss of $mCherry^+$ LET1 cells. Lower magnitude but nonetheless superior killing is also evident for lung NP63 relative to NP34 T cells. Scale bars in main figure panels indicate 1000 μm. Upper left windows are zoomed in views shown in Fig.1M relative to the larger image with the squares outlined in white dotted lines indicating their position in the original images. Scale bars in white windows indicate 100 μm. (**E)** Time-course of T cell-mediated killing of LET1 cells infected PR8 with indicated doses. mLN and lung Rg T cells were derived from RgC mice adoptively transferred with a single RgT type (dpi 7). (**F**) Intracellular NP protein expression in LET1 cells infected with the indicated dose of PR8 and then analyzed by flow cytometry. (**G, H**) Representative FACS plots (**G**) and the quantification (**H**) of NP34 ($mCherry^+$) and NP63 ($GFP^+$) T cells in lungs of mixed RgC mice 7 days after PR8 infection. Intraparenchymal lung T cells shown are identified by lack of anti-CD8α staining after *in vivo* intravenous staining in conjunction with anti-CD8β reactivity following *in vitro* staining. (**I**) Viral titer in lungs of single NP34- and 63-RgC mice (dpi 7). For **A-C, G, and H**, data are representative of four independent experiments. For **D-F, and I**, data are representative of two independent experiments and are shown as means ± SDs of six to eight mice. For all data with statistics, ****P <0.0001, ***P <0.001, *P<0.05; ns, not significant. P values were calculated by comparing slopes using linear regression analysis (**E**), paired t-test (**C and H**), or unpaired t-test (**I**).

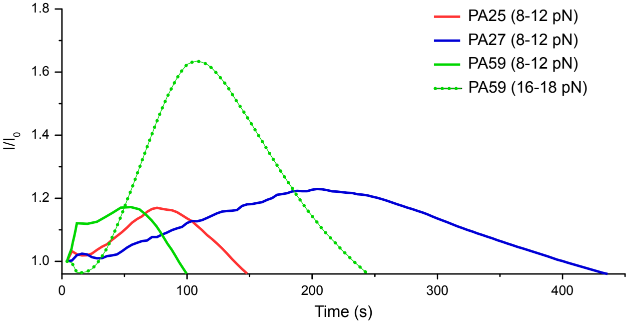

PA25 (8-12 pN)
PA27 (8-12 pN)
PA59 (8-12 pN)
PA59 (16-18 pN)
1.8
1.6
1.4
1.2
1.0
$I/I_0$
0
100
200
300
400
Time (s)

**Fig. S5. SCAR assay revealing the prolonged calcium flux in PA27 and the strong magnitude in PA59 under higher force.**

Time-course of calcium flux signal indicated as the ratio of maximum fluorescence intensity ($I_{max}$) to the initial fluorescence intensity ($I_0$) of the $Ca^{2+}$-sensitive dye for PA25-, 27-, 59-BW cells at 8-12 pN (solid lines) and PA59 at 16-18 pN (dotted line).

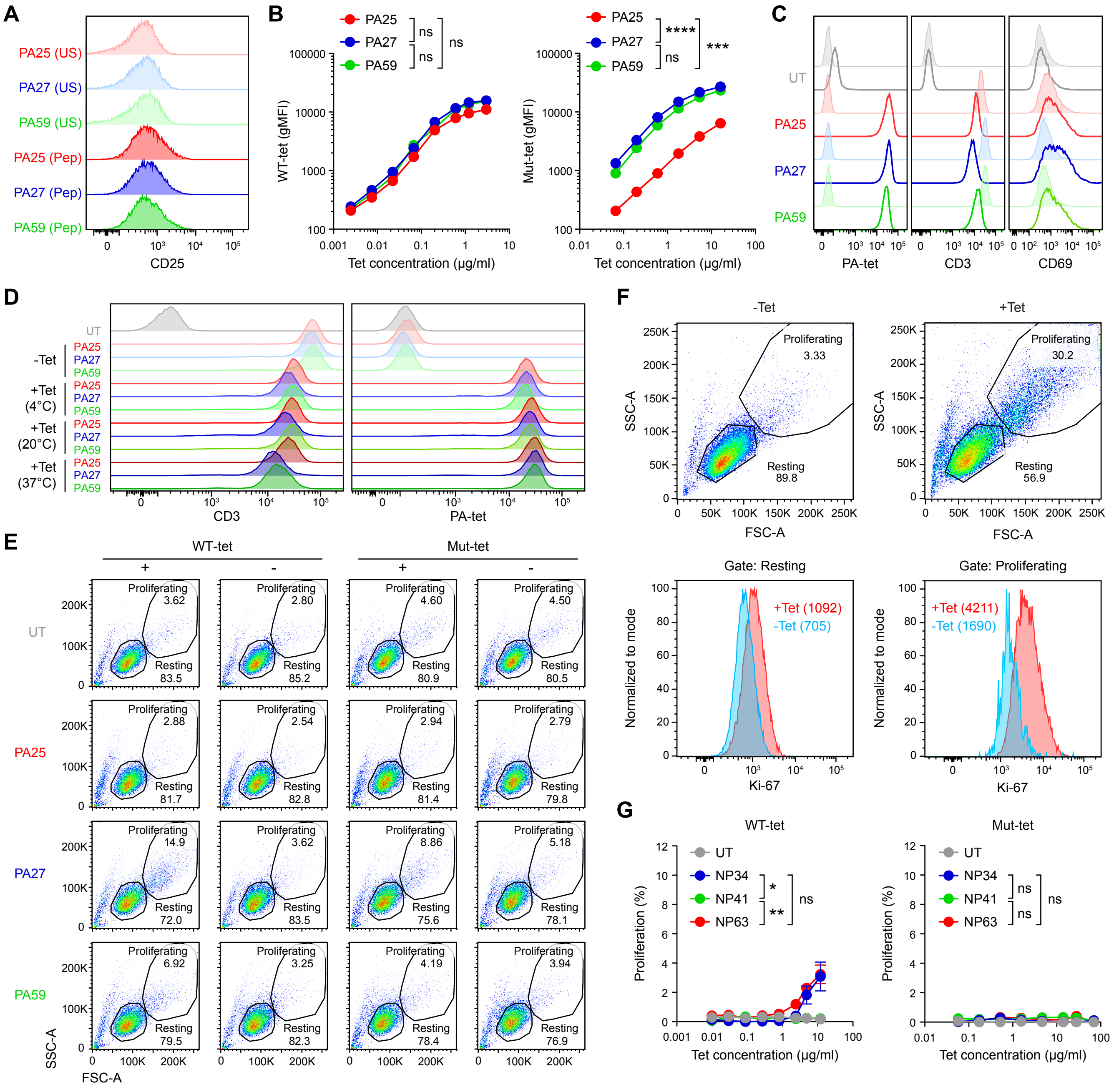

A
PA25 (US)
PA27 (US)
PA59 (US)
PA25 (Pep)
PA27 (Pep)
PA59 (Pep)
CD25
B
PA25
PA27
PA59
ns
ns
ns
WT-tet (gMFI)
Tet concentration (μg/ml)
****
***
Mut-tet (gMFI)
C
UT
PA-tet
CD3
CD69
D
-Tet
+Tet (4°C)
+Tet (20°C)
+Tet (37°C)
E
WT-tet
Mut-tet
Proliferating
Resting
PA59
SSC-A
FSC-A
F
-Tet
+Tet
Proliferating 3.33
Resting 89.8
Proliferating 30.2
Resting 56.9
Gate: Resting
Gate: Proliferating
+Tet (1092)
-Tet (705)
+Tet (4211)
-Tet (1690)
Normalized to mode
Ki-67
G
NP34
NP41
NP63
Proliferation (%)

**Fig. S6. The greatest proliferation and activation of PA TCR-expressing cells is observed for PA27 after tetramer stimulation *in vitro*.**

(**A**) CD25 expression on the indicated TCR-transduced BW cells unstimulated (US) or stimulated with 10μg/ml $PA_{224-233}$ peptide (Pep) overnight. (**B**) Tetramer binding of $PA_{224-233}$ /$D^b$ WT-tetramer (left) and CD8BS mutant-tetramer (right) for the indicated BW cell lines. Tetramer was treated for 30 minutes at 20 °C. (**C**) Representative histograms of PA-tetramer binding (left) as well as CD3 (middle) and CD69 expression (right) for PA25-, 27-, 59-, and untransduced-BW cell lines following overnight stimulation with 10 μg/mL $PA_{224-233}$ /$D^b$ WT-tetramer at 37 °C (bold lines), as shown in Fig. 2E, H, I, respectively. Shaded histograms represent control culture without tetramer addition. (**D**) CD3 expression (left) and tetramer binding (right) on indicated BW cells 30 minutes after $PA_{224-233}$ /$D^b$ WT-tetramer stimulation at indicated temperatures. (**E)** Representative FACS plots of FSC-A and SSC-A for the indicated BW cells cultured for 1 hour at 37 °C with $PA_{224-233}$ /$D^b$ WT-tetramer (3 μg/mL), its CD8BS-mutant tetramer variant (22 μg/mL), or no tetramer. The cells with larger FSC and SSC are identified as proliferating cells shown in Fig. 2J and K. (**F**) Proliferation molecule Ki67 expression in PA27-BW cells with or without $PA_{224-233}$ /$D^b$ WT-tetramer stimulation (1.2 μg/mL) for 1 hour at 37 °C. Top panels show proliferating and resting cells based on SSC-A and FSC-A scatter used for gating to determine Ki67 levels shown in the bottom row. With values in parentheses indicators gMFI. (**G**) Proliferation of indicated NP-BW cells with $NP_{366-374}$ /$D^b$ WT-tetramer (left) and CD8BS mutant-tetramer (right) stimulation. Data are normalized by subtracting the percentage of proliferating cells without stimulation from the percentage with stimulation. Data are shown as means ± SEMs of technical replicates. For, **A-C, E** and **G**, data are representative of two independent experiments. For **B** and **G**, ****P <0.0001,

***P <0.001, **P <0.01, *P<0.05; ns, not significant. P values were calculated by comparing slopes of linear regression.

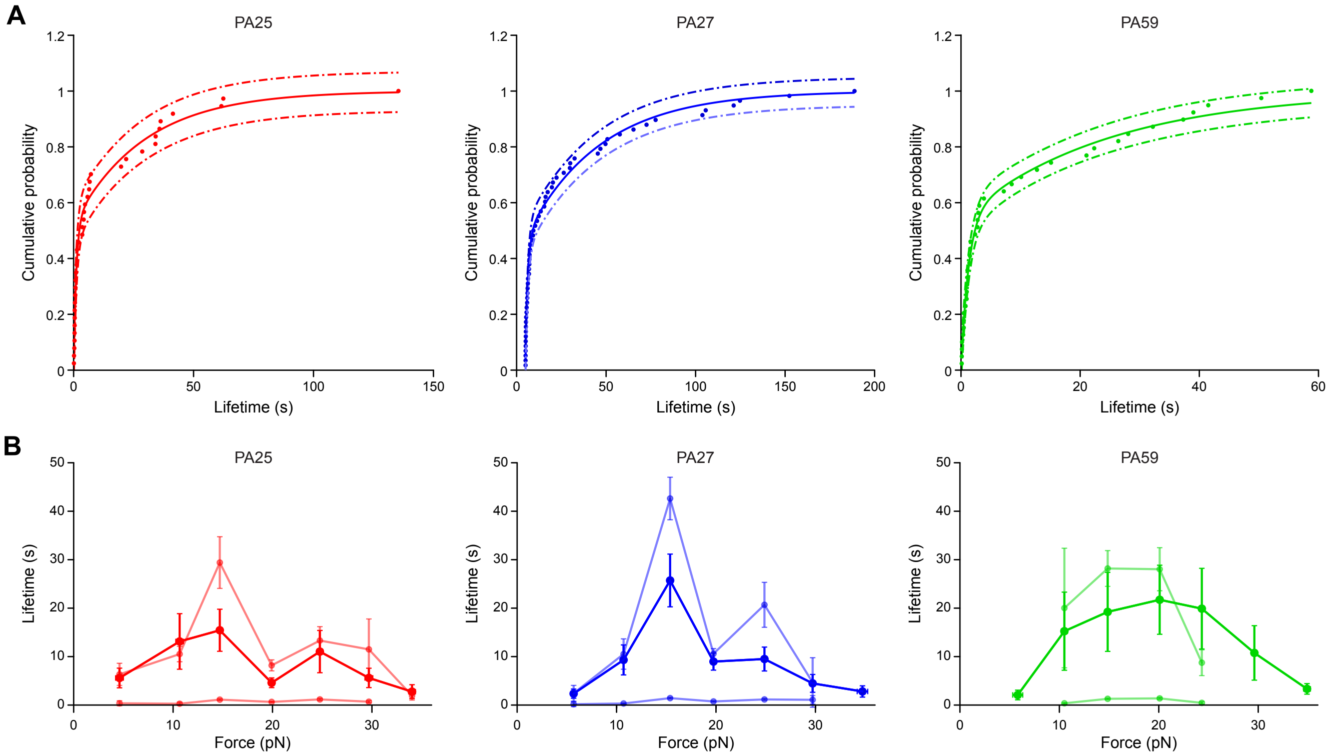

A
PA25
PA27
PA59
Cumulative probability
Lifetime (s)
B
PA25
PA27
PA59
Lifetime (s)
Force (pN)

**Fig. S7. Bin by bin cumulative probability distributions fit a double exponential with long- and short- time constants for dissociation.**

**(A)** Double exponential fit to cumulative probability distribution of lifetimes within the 15 pN bin for PA25 (red), PA27 (blue), and PA59 (green). Equation used was $y = A * \left(1 - e^{-x/t1}\right) + B * \left(1 - e^{-x/t2}\right)$. Solid lines show fit with 95% confidence intervals shown by dashed lines. **(B)** Plots comparing time constants from the double exponential fit (light colors) to the averages from the catch bond curves (dark colors) for each clone, respectively. For time constants, 95% confidence for each parameter are shown. In contrast, catch bond averages are plotted with SEM. In each PA receptor system there is an underlying baseline < 2 seconds of quick dissociation events, and a second population of long lifetime events.

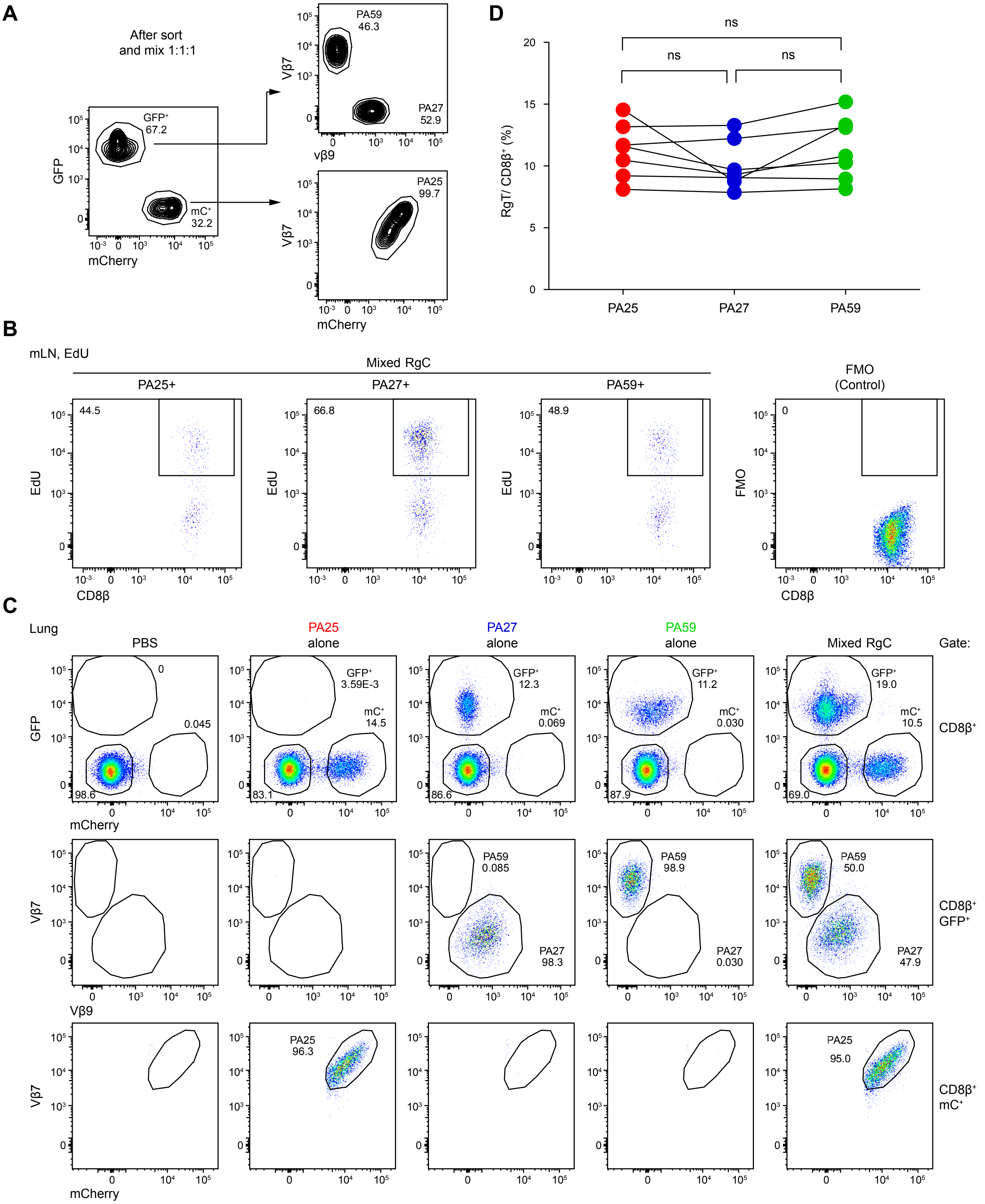
A
After sort
and mix 1:1:1
GFP⁺
67.2
mC⁺
32.2
GFP
mCherry
PA59
46.3
PA27
52.9
Vβ7
vβ9
PA25
99.7
Vβ7
mCherry
D
ns
ns
ns
RgT/ CD8β⁺ (%)
PA25
PA27
PA59
B
mLN, EdU
Mixed RgC
PA25+
PA27+
PA59+
FMO
(Control)
44.5
66.8
48.9
0
EdU
FMO
CD8β
C
Lung
PBS
PA25
alone
PA27
alone
PA59
alone
Mixed RgC
Gate:
GFP⁺
3.59E-3
mC⁺
14.5
GFP⁺
12.3
mC⁺
0.069
GFP⁺
11.2
mC⁺
0.030
GFP⁺
19.0
mC⁺
10.5
0
0.045
98.6
83.1
86.6
87.9
69.0
CD8β⁺
GFP
mCherry
PA59
0.085
PA27
98.3
PA59
98.9
PA27
0.030
PA59
50.0
PA27
47.9
CD8β⁺
GFP⁺
Vβ7
Vβ9
PA25
96.3
PA25
95.0
CD8β⁺
mC⁺
Vβ7
mCherry

**Fig. S8. Analysis of PA-specific T cells in mixed RgC mice after IAV infection.**

(**A**) Representative FACS plots of RgT cells after cell sorting and mixing PA25, 27, and 59 RgT cells at a 1:1:1 ratio. PA25 is identified as mCherry$^{+}$ V$\beta$7$^{+}$, PA27 is as GFP$^{+}$ V$\beta$9$^{+}$, PA59 is as GFP$^{+}$ V$\beta$7$^{+}$ cells. Those cells are adoptively transferred into recipient B6 mice to generate mixed RgC mice. (**B)** Representative FACS plots of EdU incorporation in PA25, PA27, and PA59 RgT cells in mediastinal LN (mLN) of mixed RgC mice 7 days after PR8 infection, as shown in Fig. 4D. (**C, D)** Representative FACS plots (**C**) of T cell populations in the lungs of mixed RgC as well as individual Rg populations or controls and their quantification (**D**) given as the % of PA25, PA27, and PA59 lung resident Rg T cells of mixed RgC mice 7 days after PR8 infection. For **B-D**, data are representative of four independent experiments. P values were calculated by paired t-test **(D).** ns, not significant.

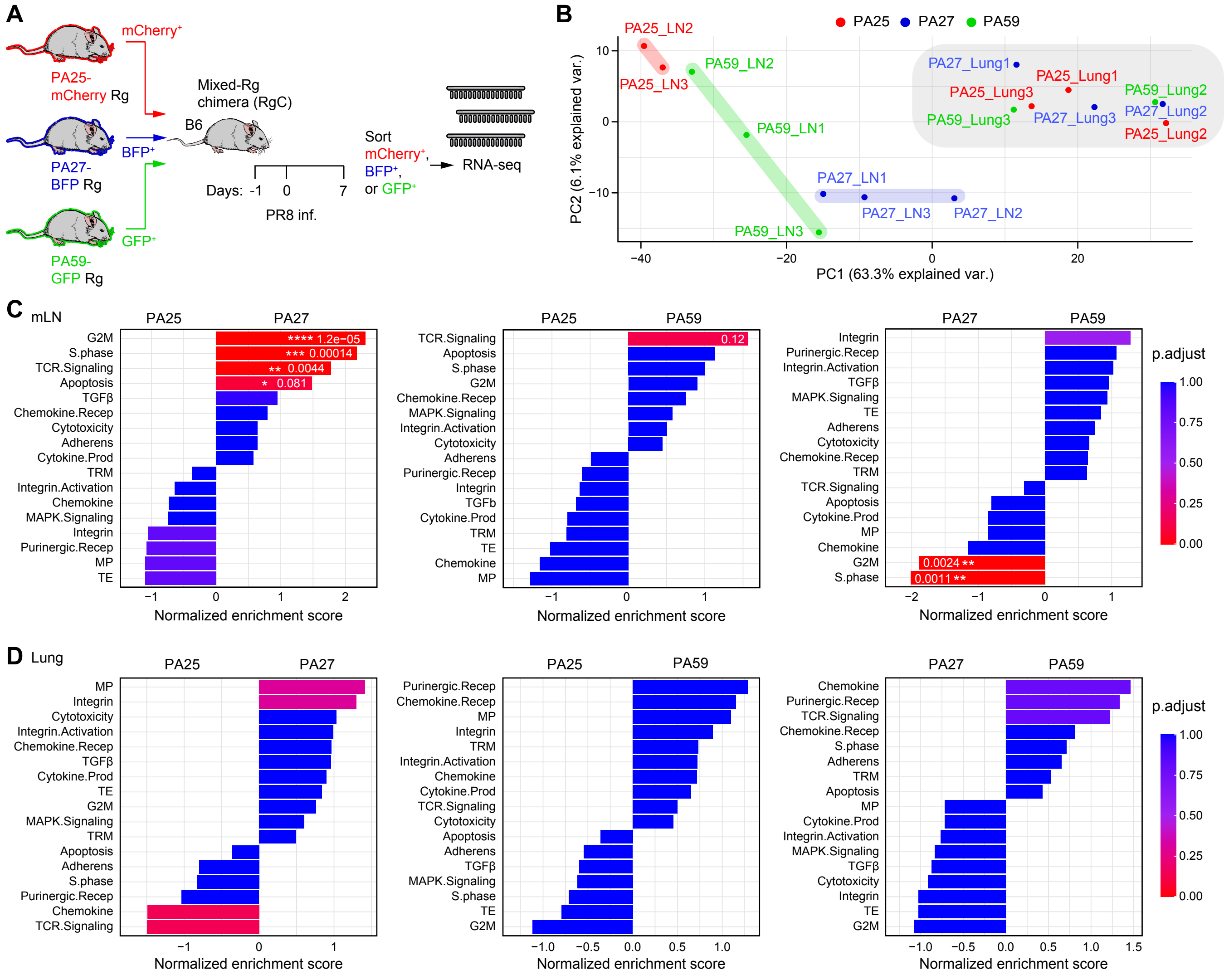
A
mCherry+
PA25-mCherry Rg
BFP+
PA27-BFP Rg
GFP+
PA59-GFP Rg
Mixed-Rg chimera (RgC)
B6
Days: -1 0 7
PR8 inf.
Sort mCherry+, BFP+, or GFP+
RNA-seq
B
PA25 PA27 PA59
PA25_LN2
PA25_LN3
PA59_LN2
PA59_LN1
PA59_LN3
PA27_LN1
PA27_LN3
PA27_LN2
PA27_Lung1
PA25_Lung1
PA25_Lung3
PA59_Lung2
PA27_Lung2
PA59_Lung3
PA27_Lung3
PA25_Lung2
PC2 (6.1% explained var.)
PC1 (63.3% explained var.)
C
mLN
PA25 PA27
G2M **** 1.2e−05
S.phase *** 0.00014
TCR.Signaling ** 0.0044
Apoptosis * 0.081
TGFβ
Chemokine.Recep
Cytotoxicity
Adherens
Cytokine.Prod
TRM
Integrin.Activation
Chemokine
MAPK.Signaling
Integrin
Purinergic.Recep
MP
TE
PA25 PA59
TCR.Signaling 0.12
Apoptosis
S.phase
G2M
Chemokine.Recep
MAPK.Signaling
Integrin.Activation
Cytotoxicity
Adherens
Purinergic.Recep
Integrin
TGFb
Cytokine.Prod
TRM
TE
Chemokine
MP
PA27 PA59
Integrin
Purinergic.Recep
Integrin.Activation
TGFβ
MAPK.Signaling
TE
Adherens
Cytotoxicity
Chemokine.Recep
TRM
TCR.Signaling
Apoptosis
Cytokine.Prod
MP
Chemokine
G2M 0.0024 **
S.phase 0.0011 **
Normalized enrichment score
p.adjust
1.00
0.75
0.50
0.25
0.00
D
Lung
PA25 PA27
MP
Integrin
Cytotoxicity
Integrin.Activation
Chemokine.Recep
TGFβ
Cytokine.Prod
TE
G2M
MAPK.Signaling
TRM
Apoptosis
Adherens
S.phase
Purinergic.Recep
Chemokine
TCR.Signaling
PA25 PA59
Purinergic.Recep
Chemokine.Recep
MP
Integrin
TRM
Integrin.Activation
Chemokine
Cytokine.Prod
TCR.Signaling
Cytotoxicity
Apoptosis
Adherens
TGFβ
MAPK.Signaling
S.phase
TE
G2M
PA27 PA59
Chemokine
Purinergic.Recep
TCR.Signaling
Chemokine.Recep
S.phase
Adherens
TRM
Apoptosis
MP
Cytokine.Prod
Integrin.Activation
MAPK.Signaling
TGFβ
Cytotoxicity
Integrin
TE
G2M
Normalized enrichment score
p.adjust

**Fig. S9. mLN PA27 T cells are strongly transcriptionally activated after IAV infection.**

**(A**) Experimental schema of a TCR labeling and sorting system for bulk RNA-seq. Different fluorescence-tagged and TCR-expressing PA-Rg T cells were sorted as fluorescence$^{+}$ CD8β$^{+}$CD44$^{-}$, mixed with at a ratio of 1:1:1, and then adoptively transferred into recipient B6 mice. These animals then were infected with PR8 24 hours post-transfer. At seven-days post-infection, Rg T cells in mLN or lungs of the mixed RgC mice were sorted as fluorescence protein$^{+}$, intravenous staining of CD8α$^{-}$, and *in vitro* staining of CD8β$^{+}$ cells and populations used to perform RNA-seq. No anti-TCR mAbs were used for this experiment. Gating for cell sorting is shown in data S3. (**B)** Principal Component Analysis (PCA) of mLN and lung samples for PA25-, PA27-, and PA59-Rg T cells. (**C, D)** Gene Set Enrichment Analysis (GSEA) based on pair-wise gene expression comparison between PA-Rg T cell samples in mLN (**C**) and lung (**D**). Statistical significance of gene set enrichment is indicated with asterisks for several thresholds of adjusted P-values (****P < 0.0001, ***P < 0.001, **P < 0.01, *P < 0.1).

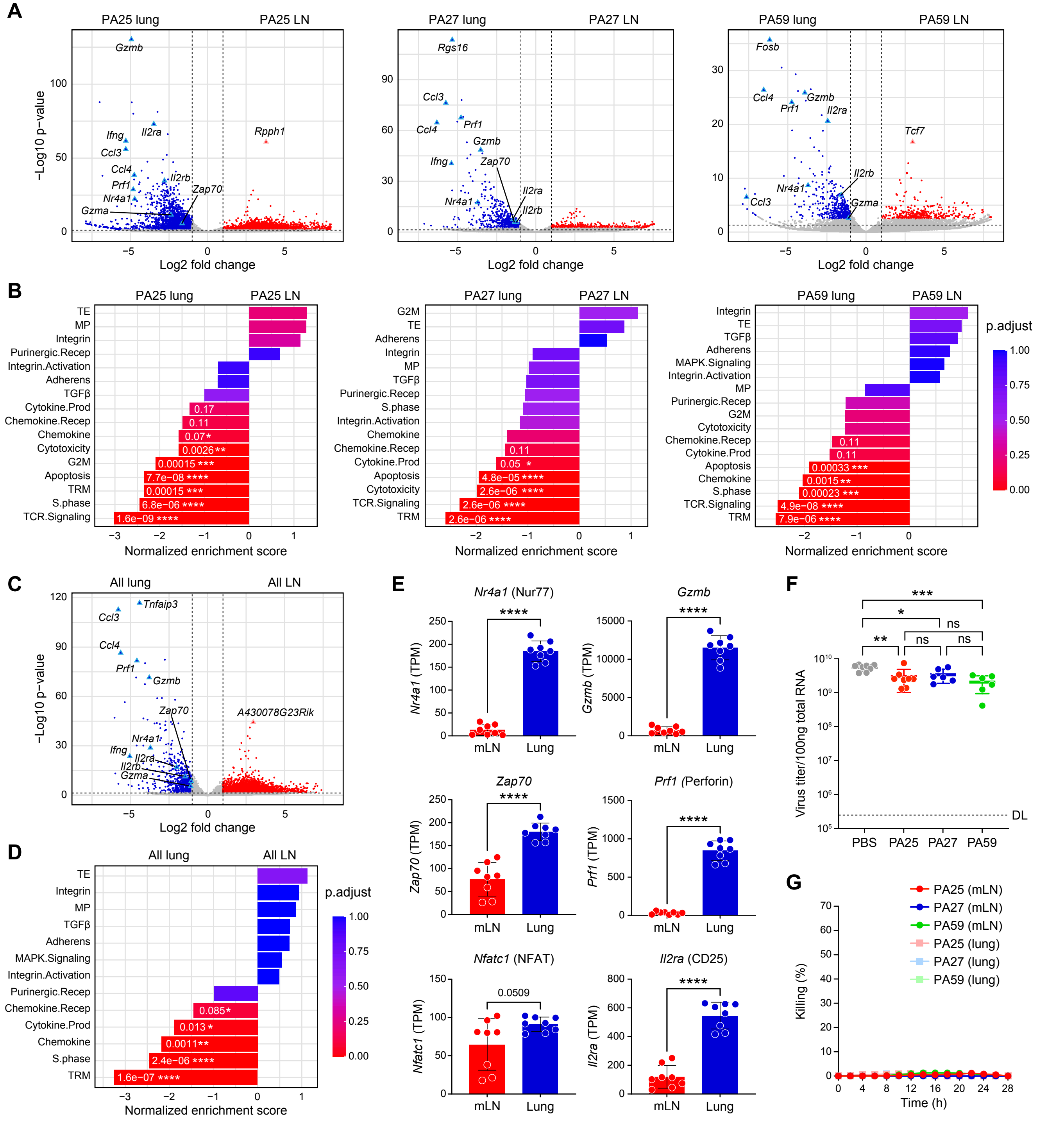
A
PA25 lung
PA25 LN
PA27 lung
PA27 LN
PA59 lung
PA59 LN
−Log10 p−value
Log2 fold change
B
Normalized enrichment score
p.adjust
C
All lung
All LN
D
E
Nr4a1 (Nur77)
Gzmb
Zap70
Prf1 (Perforin)
Nfatc1 (NFAT)
Il2ra (CD25)
mLN
Lung
0.0509
F
Virus titer/100ng total RNA
PBS
PA25
PA27
PA59
DL
ns
G
PA25 (mLN)
PA27 (mLN)
PA59 (mLN)
PA25 (lung)
PA27 (lung)
PA59 (lung)
Killing (%)
Time (h)

**Fig. S10. All lung Rg T cells significantly increase various activation genes after IAV infection.**

(**A**) Volcano plots showing differentially expressed genes (DEGs) between mLN and lung of PA25 (left), PA27 (middle), and PA59 (right). Significantly up- and down-regulated genes indicated with red and blue dots (fold-change threshold of 2 and adjusted P-value threshold of 0.05). Remarkably expressing-, effector-, cytotoxic-, and TCR signaling-genes are labeled and shown as triangles. (**B**) GSEA results based on gene expression comparison between mLN and lung of PA25 (left), PA27 (middle), and PA59 (right). Statistical significance is indicated for several thresholds of adjusted P-values with asterisks (****P < 0.0001, ***P < 0.001, **P < 0.01, *P < 0.1). (**C**,**D)** Volcano and GSEA plots similar to (**A**) and (**B**), but representing results for the comparisons of aggregated Rg T cells in mLN and lung. (**E)** Indicated gene expression in aggregated Rg T cells in mLN and lung. (**F**) Viral titer in lungs of single PA25-, PA27-, PA59-RgC mice (dpi 7) determined by real-time PCR. Control mice were injected PBS without Rg T cells when RgC mice were generated. For **E** and **F**, data are shown as means ± SDs of eight samples (**E**) or six to eight mice (F). ****P <0.0001, ***P <0.001, **P <0.01, *P<0.05; ns, not significant. P values were calculated by unpaired t-test. (**G**) Time-course of T cell-mediated killing of LET1 cells infected with PR8 at a dose of $4x10^8$ $EID_5$. mLN and lung Rg T cells were derived from RgC mice adoptively transferred with the indicated RgT type (dpi 7).

**Movie S1.**

$Ca^{2+}$ flux of PA25 T cell triggered efficiently under optimal tangential shear force (8-12 pN) but not the other forces (force outside the optimal region). The arrow indicates the direction of the force. The time in the upper left indicates minutes: seconds.

**Movie S2.**

$Ca^{2+}$ flux of PA27 T cell triggered efficiently under optimal tangential shear force (8-12 pN) but not the other forces (force outside the optimal region). The arrow indicates the direction of the force. The time in the upper left indicates minutes: seconds.

**Movie S3.**

$Ca^{2+}$ flux of PA59 T cell triggered efficiently under optimal tangential shear force (16-18 pN) but not the other forces (force outside the optimal region). Note that a weak $Ca^{2+}$ flux of PA59 T cell can be triggered under force ranging from 8 to 12 pN. The arrow indicates the direction of the force. The time in the upper left indicates minutes: seconds.

**Data S1. (separate file)**

TRV, TRJ, and CDR3 sequence for $NP_{366-374}/D^b$- and $PA_{224-233}/D^b$- specific TCRs identified by single-cell RNA-seq.

**Data S2. (separate file)**

Uncropped gels for ERK and pERK expressions of NP- (page 1) and PA-BW cells (page 2).

**Data S3. (separate file)**

Gating strategy for PA-Rg T cell sorting without utilizing anti-TCR mAbs for RNA-seq. Page 1 shows mLN cells and page 2 shows resident lung cells from 3 mixed RgC mice and control mice.

**Data S4. (separate file)**

Gene expression in mLN and lung PA25-, 27-, and 59-Rg T cells after IAV infection (dpi 7) obtained by RNA-seq. The expression level is shown by TPM (Transcript Per Million). The samples filled with gray, PA25_LN_m01 and PA59_Lung_m01 were excluded for further analysis due to their low quality of RNA.

**Data S5. (separate file)**

Antibodies and tetramers used in this study.

**Data S6. (separate file)**

The primers and detailed conditions used for single-cell RT-PCR.

**Data S7. (separate file)**

The sources of gene sets used for Gene Set Enrichment Analysis (GSEA)  analysis.
