## Supplementary figures and images for "Parsing digital or analogue TCR performance through piconewton forces"

### Data S2

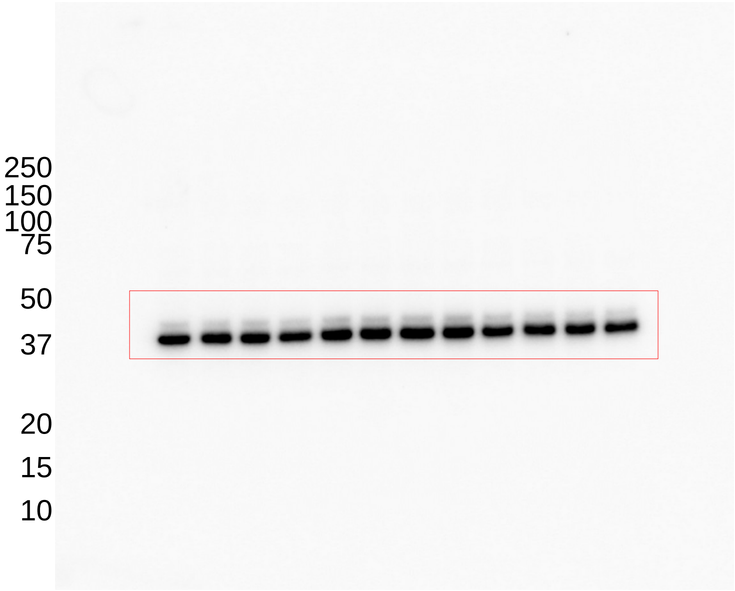


ERK: NP TCR

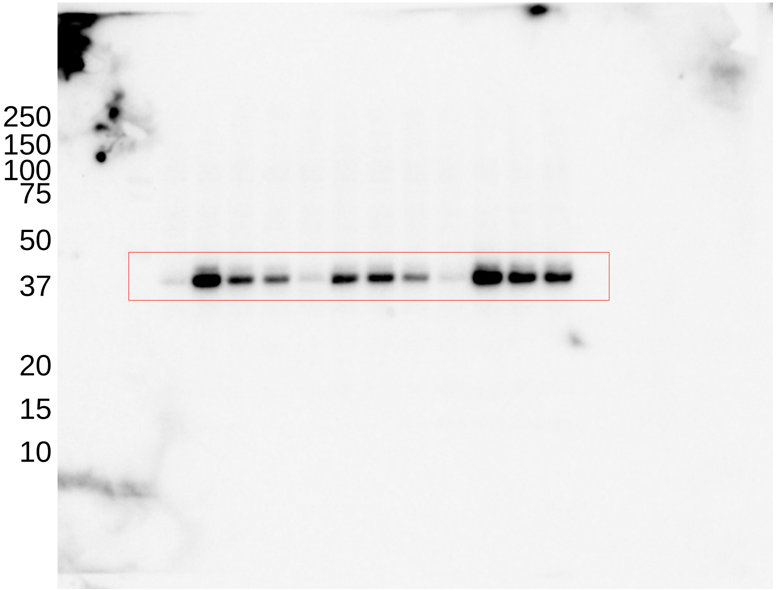


Phosphorylated ERK: NP TCR

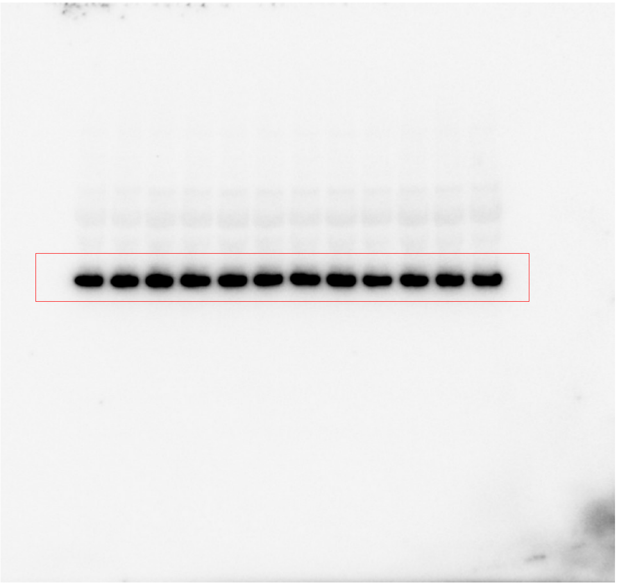

ERK: PA TCR

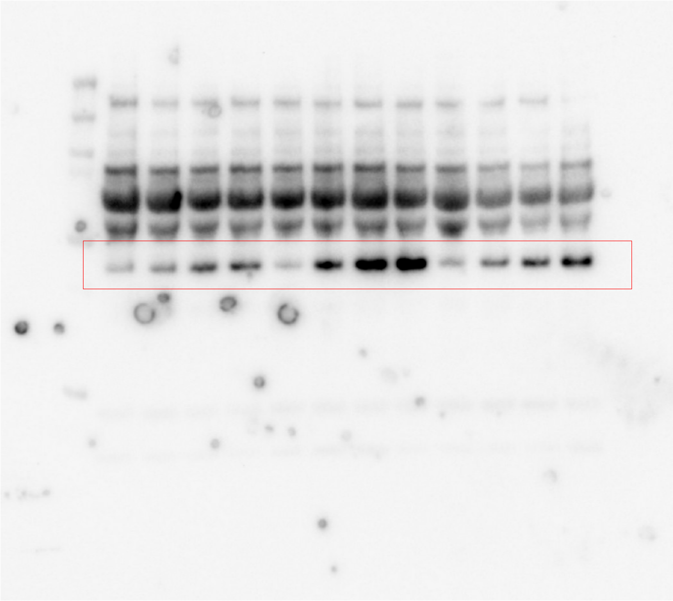

Phosphorylated ERK: PA TCR

### Data S3

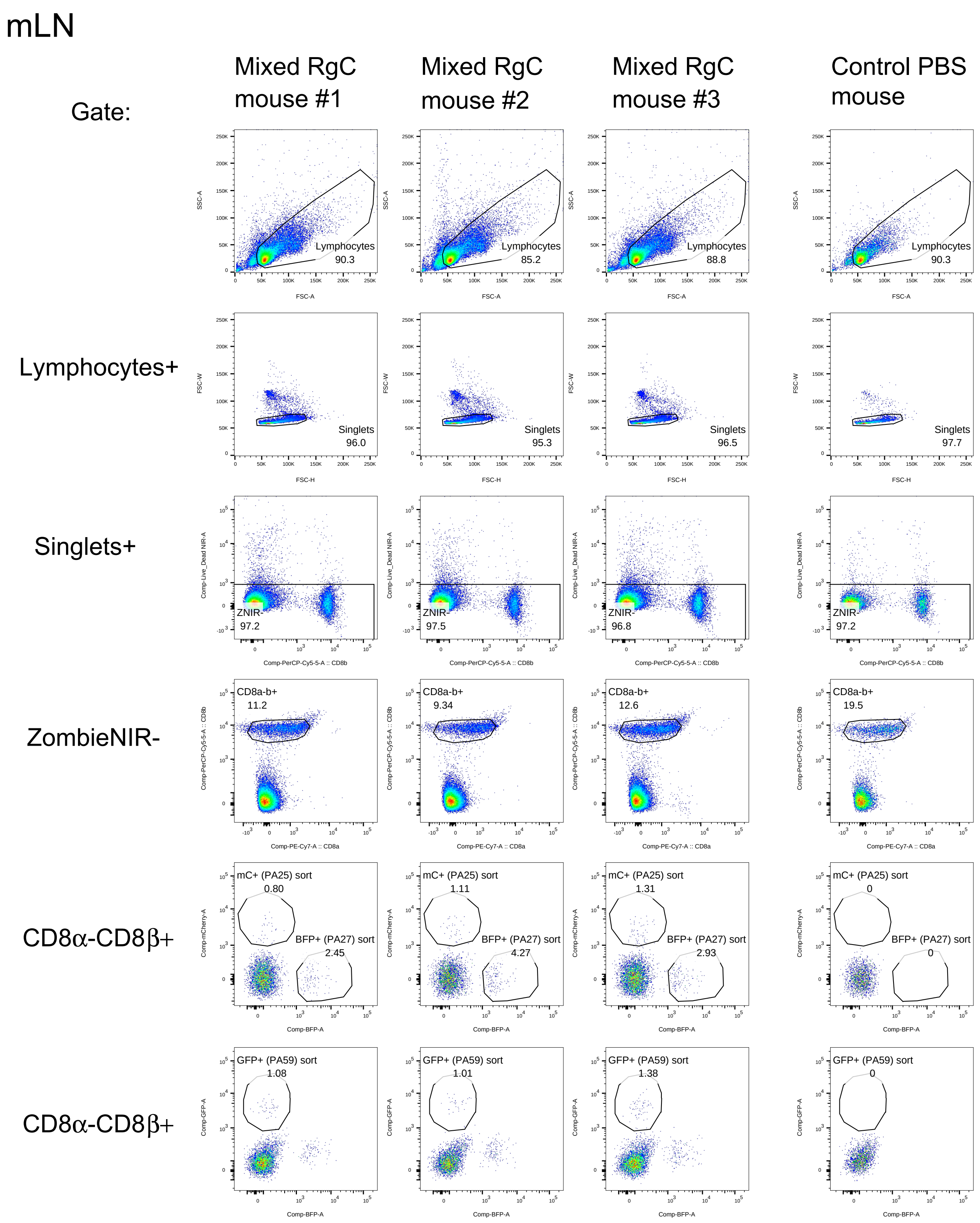

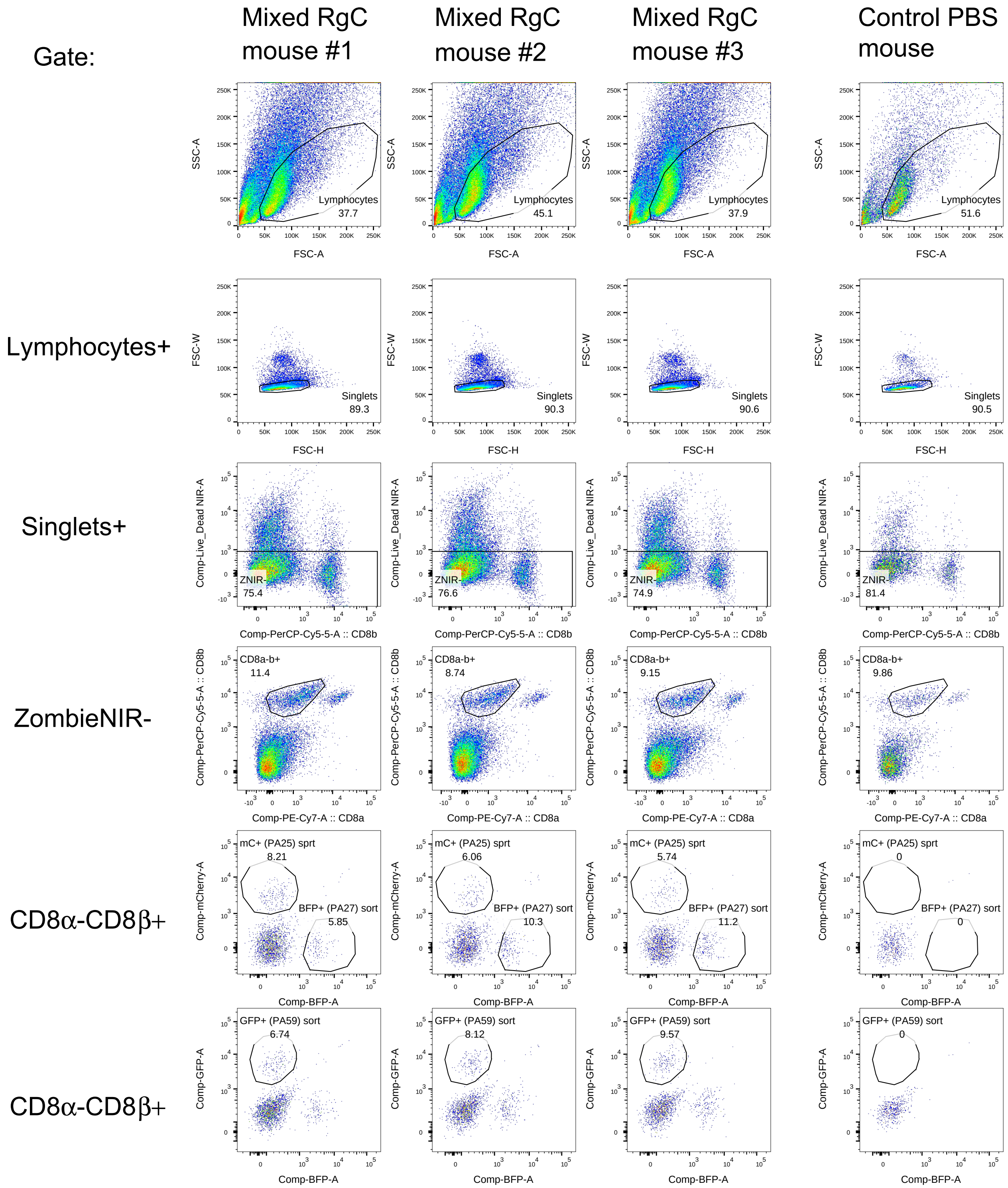
