## Supplementary material for "Parsing digital or analogue TCR performance through piconewton forces": Data S7

### Name of gene set
Exact name on database
Database_Category_Systematic name
URL
Note

### Cell adhesion
GOBP_CELL_CELL_ADHESION
GSEA_C5_M12087
https://www.gsea-msigdb.org/gsea/msigdb/cards/GOBP_CELL_CELL_ADHESION.html
Human HLA genes were excluded from the analysis.

### Integrin
REACTOME_INTEGRIN_CELL_SURFACE_INTERACTIONS
GSEA_ C2_ M16441
https://www.gsea-msigdb.org/gsea/msigdb/cards/REACTOME_INTEGRIN_CELL_SURFACE_INTERACTIONS.html

### Integrin activation
GOBP_INTEGRIN_ACTIVATION
GSEA_ C5_M23217
https://www.gsea-msigdb.org/gsea/msigdb/cards/GOBP_INTEGRIN_ACTIVATION.html

### Cytotoxicity
GOBP_LEUKOCYTE_MEDIATED_CYTOTOXICITY
GSEA_ C5_M11242
https://www.gsea-msigdb.org/gsea/msigdb/cards/GOBP_LEUKOCYTE_MEDIATED_CYTOTOXICITY.html
Human HLA genes were excluded from the analysis.

### Cytokine production
GOBP_POSITIVE_REGULATION_OF_T_CELL_CYTOKINE_PRODUCTION
GSEA_ C5_M13565
https://www.gsea-msigdb.org/gsea/msigdb/cards/GOBP_POSITIVE_REGULATION_OF_T_CELL_CYTOKINE_PRODUCTION.html
HLA genes were excluded from the analysis.

### Chemokine
GOMF_CCR_CHEMOKINE_RECEPTOR_BINDING
GSEA_ C5_M18725
https://www.gsea-msigdb.org/gsea/msigdb/cards/GOMF_CCR_CHEMOKINE_RECEPTOR_BINDING.html
Cytoplasm genes, JAK1, NARS1, STAT1 were excluded from the analysis.

### Chemokine receptor
GOMF_CHEMOKINE_BINDING

GSEA_ C5_M18707
https://www.gsea-msigdb.org/gsea/msigdb/cards/GOMF_CHEMOKINE_BINDING.html
HMGB1 was excluded from the analysis.

### Purinergic receptor

GOMF_G_PROTEIN_COUPLED_PURINERGIC_NUCLEOTIDE_RECEPTOR_ACTIVITY
GSEA_ C5_M34434
https://www.gsea-msigdb.org/gsea/msigdb/cards/GOMF_G_PROTEIN_COUPLED_PURINERGIC_NUCLEOTIDE_RECEPTOR_ACTIVITY.html

### Apoptosis

KEGG_APOPTOSIS
GSEA_ C2_ M8492
https://www.gsea-msigdb.org/gsea/msigdb/cards/KEGG_APOPTOSIS.html

### Adherens_Junction pathway

Adherens junction
KEGG_map04520
https://www.genome.jp/dbget-bin/www_bget?pathway:map04520

### TCR signaling pathway

T cell receptor signaling pathway
KEGG_map04660
https://www.genome.jp/dbget-bin/www_bget?pathway:map04660
CD4 was excluded from the analysis.

### TGFb signaling pathway

TGF-beta signaling pathway
KEGG_map04350
https://www.genome.jp/dbget-bin/www_bget?pathway:map04350

### MAPK signaling pathway

MAPK signaling pathway
KEGG_map04010
https://www.genome.jp/dbget-bin/www_bget?pathway:map04010
